# Colocalization of Protein and microRNA Markers Reveals Unique Extracellular Vesicle Sub-Populations for Early Cancer Detection

**DOI:** 10.1101/2023.04.17.536958

**Authors:** Zongbo Li, Kaizhu Guo, Ziting Gao, Junyi Chen, Zuyang Ye, Shizhen Emily Wang, Yadong Yin, Wenwan Zhong

**Author notes:** Corresponding Author Wenwan Zhong – Professor of Chemistry, University of California-Riverside, Riverside, CA 92521, USA. orcid.org/0000-0002-3317-3464. Authors contributed equally to the work (Zongbo Li, Kaizhu Guo, Ziting Gao).

## Abstract

Extracellular vesicles (EVs) play important roles in cell-cell communication but they are highly heterogeneous, and each vesicle has dimensions smaller than 200 nm thus encapsulates very limited amounts of cargos. We report the technique of NanOstirBar (NOB)-EnabLed Single Particle Analysis (NOBEL-SPA) that utilizes NOBs, which are superparamagnetic nanorods easily handled by a magnet or a rotating magnetic field, to act as isolated “islands” for EV immobilization and cargo confinement. NOBEL-SPA permits rapid inspection of single EV with high confidence by confocal fluorescence microscopy, and can assess the colocalization of selected protein/microRNA (miRNA) pairs in the EVs produced by various cell lines or present in clinical sera samples. Specific EV sub-populations marked by the colocalization of unique protein and miRNA combinations have been revealed by the present work, which can differentiate the EVs by their cells or origin, as well as to detect early-stage breast cancer (BC). We believe NOBEL-SPA can be expanded to analyze the co-localization of other types of cargo molecules, and will be a powerful tool to study EV cargo loading and functions under different physiological conditions, and help discover distinct EV subgroups valuable in clinical examination and therapeutics development.

---

Extracellular Vesicles (EVs) secreted by cells can mediate cell-cell communication,^1–5^ and are present in all biological fluids,^6, 7^ easily accessible with minimal invasion.^8^ They are classified into different subtypes;^9–15^ and the two smaller (diameter ∼40-250 nm) subtypes called exosomes and microvesicles have unique biogenesis pathways and diverse types of cargo molecules,^1–3, 5, 16^ attracting great attention in biomedical research. EVs have been associated with immune responses, viral pathogenicity, cancer progression, and cardiovascular or central nervous system–related diseases,^17–20^ supporting their high potential as diagnostic and therapeutic tools. However, it is very challenging to identify the disease-related EVs. EVs are highly heterogeneous, different in their sizes, contents, cells of origin, biogenesis pathways, and functional impacts on recipient cells.^21–23^ While the total EV concentration in the peripheral circulation can reach 10^9^ vesicles/mL,^24^ the unique EV sub-populations that carry out specific disease-related functions, or are derived from cells undergoing pathological transition^21, 25^ could be at very low abundance during the early development stage.^26^ It has been projected mathematically that, for bulk detection, it would need techniques sensitive enough to accommodate an EV input of ∼ 100 EV particles/mL in order to detect those released by small human tumors (< 1 cm^3^), which are curable if caught early.^27^ Such a sensitivity requirement is very difficult if not impossible to be met by the conventional methods of ELISA, Western Blotting (WB), and bead-based flow cytometry (FCM). In addition, bulk analysis only produces the ensemble average of the varying signals from a swarm of heterogeneous EVs, and very likely miss the signals from the distinct sub-populations present at trace levels.

To simultaneously overcome the heterogeneity and sensitivity issues in EV analysis, pioneering works analyzed single EVs using high-resolution FCM,^28–31^ super resolution microscopy,^32, 33^ and droplet-based NGS.^34^ Confocal or total internal reflection fluorescence microscopy (CFM and TIRFM) has also come to the spot light of single EV analysis because of their relatively lower cost in instrumentation and less complex operation, while offering direct visualization of the vesicles.^35, 36^ As listed in **Table S1,** the most state-of-the-art developments using fluorescence microscopy detected either proteins or nuclei acids on single EVs, confirming the presence of EV sub-populations bearing different phenotypic features and their premises in marking disease development.^37–41^ Still, single EV analysis is very difficult, owing to their extremely small sizes and the low amounts of cargos enclosed in each EV. While impressive detection performance has been obtained with the pioneering developments, assay turn-around time, limit of detection (LOD), and sample consumption are yet to be improved to meet the needs in early detection and frequent disease monitoring.^37–41^ Single EV capture was only achieved with specially designed surface features fabricated on microfluidic devices; or by controlling a large bead-to-EV molar ratio, which is not easy to do when testing unknown samples. Analyzing multiple types of cargos in single EVs with comparable sensitivity as protein detection has not yet been achieved either, which is significant for better understanding of EV cargo selection and EV functions.^42^

Herein, we report the technique of NanOstirBar (NOB)-EnabLed Single Particle Analysis (NOBEL-SPA) that employs the multifunctional NOBs to enable detection of both the protein and microRNA (miRNA) cargos on single EVs relying on the unique structure of NOBs. The NOBs not only act as regular magnetic particles to facilitate easy handling, but also can spin freely in the rotating magnetic field to speed up molecular diffusion to the NOB surface and prompt rapid target binding and thorough removal of the non-specifically adsorbed molecules. Moreover, each NOB has comparable dimensions as the single EV. Thus, it can easily realize one-NOB-one-vesicle during EV capture and then act as an isolated “island” for immobilization of the individual EV and its cargo molecules, preventing EV aggregation and diffusion of the intra-vesicular molecules once the membrane structure is destroyed. The few marker molecules concentrated on each NOB can be illuminated by the DNA nanoflowers (**DNF**) grown from rolling circle amplification (**RCA**),^43^ making the single EV carrying specific markers easily detected with high confidence by diffraction-limited confocal fluorescence microscopy. The effectiveness of NOBEL-SPA was demonstrated through the detection of both tumor protein and miRNA markers in the exosomes produced by various tumor cell lines or present in the sera samples collected from breast cancer (**BC**) patients. Distinct EV sub-populations defined by the colocalization of specific tumor miRNA and protein markers were found to be able to differentiate EVs by their cells of origin and to differentiate Stage I BC patients from healthy controls. We believe NOBEL-SPA can be applied to study other sub-micron biological particles, help gain more understanding of their cargo loading and functions, and identify specific sub-populations carrying unique features.

## Results and Discussion

### NOBs to facilitate single EV counting and yield efficient EV capture

Our previous work^44, 45^ have employed fluorescent DNA nanoflowers (DNF) to amplify the size and signals from surface proteins of individual EVs so that they can be easily seen by a conventional flow cytometer or a diffraction-limited confocal microscope. DNF construction is triggered by the binding between a protein marker and an aptamer probe^44^ or a primer-conjugated antibody.^45^ The binding initiates rolling circle amplification (RCA) and grows a long ssDNA product, the sequence of which permits self-hybridization to fold into a nanostructure. We proved that, with each DNF labeled by numerous fluorophores, detection of single EVs carrying protein markers, like HER2, EGFR, CD44, etc., can be easily done by CFM with an input ∼100 EV particles (P)/µL.^45^ But EV capture on the flat glass surface by antibodies, similar to other pioneering works, is slow and inefficient due to limitation in surface diffusion. Additionally, EVs aggregate easily, preventing accurate recognition of individual vesicles.

In the present work, we discovered that the superparamagnetic, silica-coated nanorods^46–48^ can be used to enable single EV analysis (**Figure 1a**). While the NOBs are small (< 200 nm) and can be homogeneously dispersed in aqueous solutions, complete pull-down from a 1.5-mL solution by a magnet can be done in 10 seconds. They also can spin in solution if driven by a low-gradient rotating magnetic field generated by a stirrer plate. We synthesized the NOBs with the dimension of ∼150 nm × 50 nm and modified their silica surface with carboxyl groups. Then the NOBs were conjugated with the mixture of antibodies against the exosomal markers of CD63/CD9/CD81 to specifically capture exosomes (**Fig. 1b and S1**) from a standard EV sample purchased commercially. We stained the NOB-bound exosomes with the membrane-bound dye DiB (λ_ex_353 nm/λ_em_ 442 nm) and evaluated the total number of vesicles detected by CFM after various capture durations.

**Figure 1.**
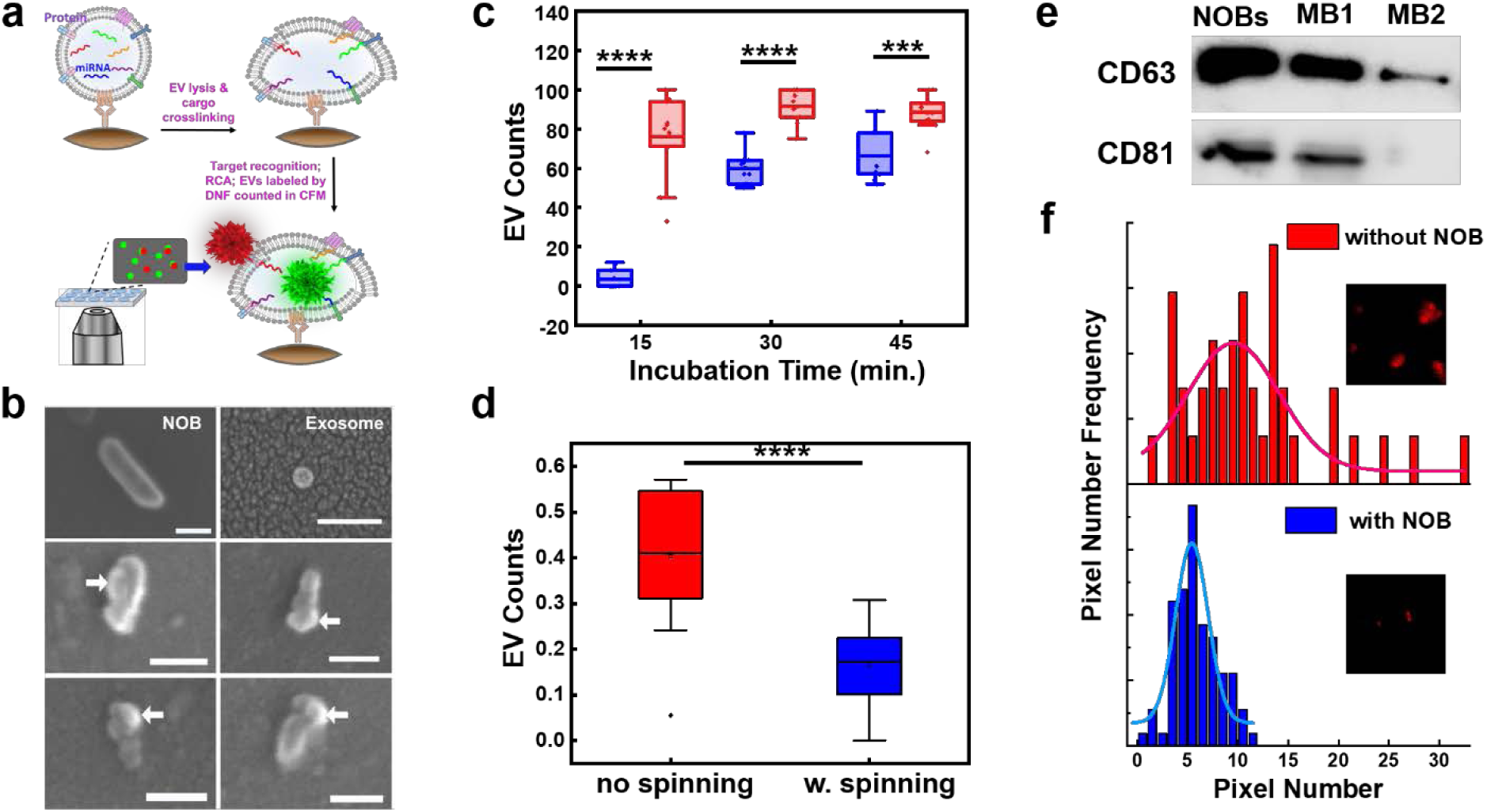
NOB to facilitate single vesicle analysis. **a)** Scheme of NOBEL-SPA. **b)** SEM images of the individual NOB, exosome, and the antibody-modified NOB with the bound exosome (pointed by the arrows). Scale bar: 100 nm. **c-d)** Box plots for the counts of DiB-stained exosomes captured by c) the antibody-conjugated NOBs under no-spinning or spinning condition; and d) the no-antibody conjugated NOBs after washing with or without spinning the NOBs. The box is determined by the 25th and 75th percentiles, with the line in the box representing the median value, and the whiskers determined by the 5th and 95th percentiles. **e)** WB results for detection of CD63 and CD81 in the exosomes pulled down by the NOBs, or the spherical magnetic beads (MBs) with diameters around 1 (MB1) or 0.22 µm (MB2), after 30-min incubation. **f)** Histograms for the pixel numbers of the fluorescent particles detected after exosome lysis and cargo crosslinking, with the exosomes captured on the glass slide surface (no NOB) or by the NOBs (inset images: 10 µm × 10 µm). ***: p < 0.001; ***: p < 0.0001.

We found that, with spinning, the NOBs took only 15 min to reach the capture plateau, with the maximum number of bound exosomes ∼20% higher than those obtained from 45 min incubation without spinning (**Fig. 1c**). In addition, spinning the NOBs carrying no antibodies during wash produced at least 2 times lower background signals compared to wash without spinning (**Fig. 1d**). Moreover, more exosomes were isolated by the NOBs than the 1- or 0.22-µm spherical magnetic microbeads (MB), as confirmed by testing the exosomal markers of CD63 and CD81 pulled down by the NOBs and the MBs with Western Blot (**Fig. 1e**). Quantification of the remaining exosomal proteins in the supernatant by ELISA also confirmed that > 85% of the exosomes were pulled down by the NOBs, while ∼ 60% were captured by the MBs (**Fig. S2**). These results well support that, the spinning action of the NOBs can promote molecular diffusion and thus improve the efficiency of target capture and impurity removal.

### NOBs to simplify miRNA detection in single vesicle

Most importantly, the NOBs can simplify single EV counting and facilitate analysis of the intravesicular cargos like microRNAs (miRNAs). Since the NOBs have the dimensions comparable to those of the EVs, each NOB can capture only few EVs due to space hindrance. We can further limit the number of EVs captured per NOB by controlling the amount of antibody (anti-CD63/CD9/CD81) loaded on each NOB. When using a molar ratio of 1 : 10 for NOB : antibody in the conjugation, either no or just 1 exosome was found on the NOB when viewed by SEM (**Fig. 1b**). With the individual EVs spatially confined and separated from each other to minimize EV aggregation, counting the single EVs illuminated by the fluorescent DNF in CFM is highly simplified. The NOBs also provide the solid support for fixing the EVs and their cargos upon breaking down the EV membrane structures by detergents, allowing analysis of the intravesicular miRNAs (**Fig. 1a**). We lysed the exosomes bound to the NOBs by a 10-min treatment of 4% paraformaldehyde (PFA);^49^ and used 1-ethyl-3-(3-dimethylaminopropyl) carbodiimide (EDC) to crosslink the 5’-phosphate of RNA^50^ to the EV proteins and antibodies on the NOBs (**Figure S3**). In this way, EV cargos would not diffuse to the surrounding area and maintain their initially high concentrations inside each EV. The exposed EV miRNAs can then hybridize with a hairpin probe to release the primer region for binding with the circular template that initiates RCA and grow the fluorescent DNF (**Figure S4a, and probe designs in Fig. S4d and S6d**). The DNF-labeled EV supported by a NOB can be imaged as a bright fluorescent particle. The sizes of the fluorescence particles resulted from the NOB-captured exosomes were much smaller (occupying only 4-8 imaging pixels) and more homogeneous (showing a narrower distribution profile of the pixel numbers per particle) compared to those not supported by the NOBs (**Fig. 1f**). A size of 4-8 pixels under our imaging condition is equivalent to a dimension of ∼ 250-300 nm, matching well with that of the single exosomes labeled with the DNS as found in our previous works.^44, 45^

Detection of two potential tumor miRNA markers, miR-155 and miR-122, on single EVs was tested in the purchased exosome standards. Good detection specificity using the hairpin probes was confirmed: only the target miRNA (miR-155 or miR-122) yielded positive RCA reaction, but not the ssRNA with small sequence variations nor other miRNAs (**Fig. S4 and S6**). We stained the exosomes with DiB; and labeled the DNF grown upon recognition of miR-155 or miR-122 with Alexa633 (λ_ex_ 621 nm/ λ_em_ 639 nm) (**Fig. 2a**). The number of the stained exosomes, **P_EV_**, and that of the DNF-labeled exosomes, **P_miRNA_**, detected by CFM were both linearly proportional to the input exosome concentration in the range of 200 – 10^5^ EV particles/µL (P/µL) (sample volume = 10 µL) when plotted in the Log scales (solid lines in **Fig. 2b; Fig. S5 and S6**). Using the 3σ method, we calculated the LOD for miR-155 and miR-122 being 3.1 P/µL and 18 P/µL, respectively. The LOD differences reflect the differential loading of these two miRNAs in this exosome sample: as calculated from the miRNA quantity obtained from RT-PCR and the exosome counts measured by Nanoparticle Tracking Analysis (NTA), at least 3 copies of miR-155 were found in each exosome; but only ∼2 copies of miR-122 were present in ∼100 exosomes (**Figure S8d**). Using P_EV_ for LOD calculation yielded a value of 4 P/µL (**Figure S5**), comparable to that of miR-155 and confirming the high abundance of miR-155. These LODs are lower than what we previously reported for EV surface protein^45^ and also that reported by others for analysis of miRNAs in single EVs using lipid vesicle fusion.^40, 51^ The improvement could be owing to the high EV capture efficiency and the effective preservation of the EV cargos on NOB.

**Figure 2.**
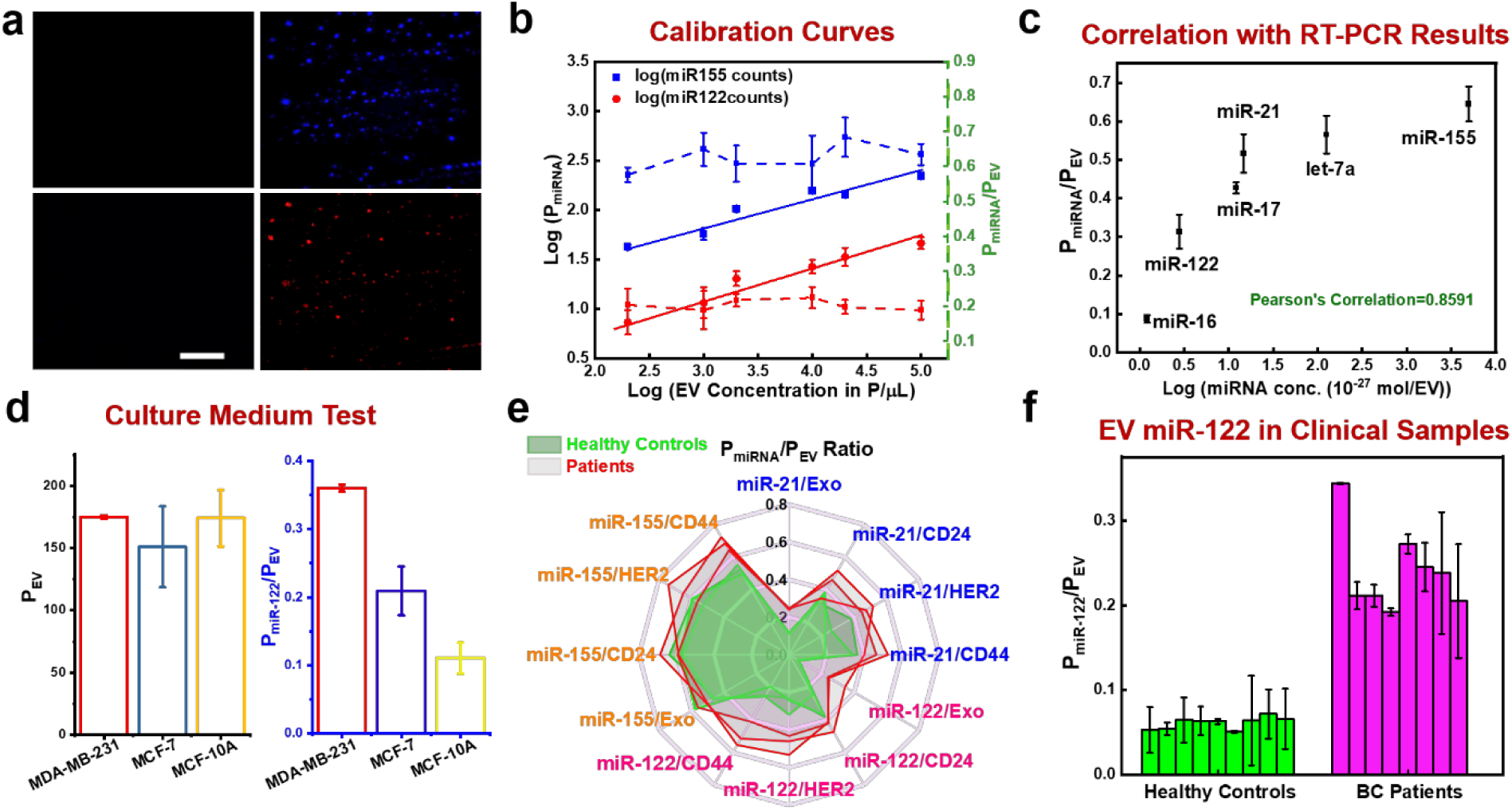
NOBEL-SPA for analysis of EV miRNAs associated with individual EVs. **a)** Representative images of the DiB-stained exosomes captured by NOBs (top panel) and illuminated by the DNF labels for miR-155 detection (bottom panel) collected with no (left panel) or 2 × 10^5^ P/µL (right panel) exosomes. Scale bar: 20 µm. **b)** Calibration curves (solid lines) for miR-155 and miR-122 obtained by plotting the Log of the PmiRNA counts vs. the Log of the input EV concentration; and the ratios of P_miRNA_/P_EV_ (dash lines) detected at various EV concentrations. c) The ratio of P_miRNA_/P_EV_ for various miRNAs positively correlated with the Log (miRNA concentration) detected by RT-PCR and NTA. d) P_EV_ and P_miRNA_/P_EV_ detected in the culture medium from MCF-10A, MCF-7, and MDA-MB-231 cells after 48 hrs EV harvest. e) Spider web plot comparing the ratios of P_miRNA_/P_EV_ obtained in the sera of 3 BC patients (red) and 3 healthy controls (green), with the EVs captured via recognition of the surface protein of CD63, HER2, CD44, or CD24, and respectively detecting the EV-associated miR-155, miR-21, or miR-122. f) The ratio of P_miRNA_/P_EV_ found in the sera of 8 BC patients (magenta columns) and 9 healthy controls (green columns), with the EVs captured by the anti-exosomal markers NOBs and detecting the EV-associated miR-122. All particle counts and counts ratio were the average values taken from two repeat measurements per sample. In each repeat, the particle counts were the total value taken from 10 images, and the ratio was the average ratio from 10 images.

Interestingly, we found that, in the same exosome sample, the **P_miRNA_/P_EV_** ratio for the same miRNA did not change significantly with different EV inputs (dash lines in **Fig. 2b**), but were distinct between two miRNAs: for miR-155 it was > 60%, but for the low-abundant miR-122 it was only ∼20%. We then designed specific stem-loop probes for various miRNAs (**Figure S7-8**), and confirmed that indeed the P_miRNA_/P_EV_ ratio varied among different miRNAs, and was positively correlated with the Log (miRNA moles per particle) quantified by RT-PCR and NTA (R^2^ = 0.8591) (**Fig. 2c**). These results prove that, the P_miRNA_/P_EV_ ratio is reflective to the abundance of miRNAs in the EV samples. By staining the EVs to count the total EVs captured by the NOBs, NOBEL-SPA can accurately compare the miRNA expression levels in different EV samples without the need to quantify the total EVs or total EV RNAs. Detection of each miRNA marker by our method requires an EV input < 2,000 particles, but > 10^7^ particles are needed by RT-PCR. Our method can work directly with unprocessed samples without RNA extraction, thus is much more efficient and faster.

### Differential loading of miRNAs in EVs from different sources

The high sensitivity and simplicity of NOBEL-SPA permit quick assessment of EV production from cells and the enclosed miRNA contents. Five µL culture medium of MCF-10A (non-tumor), MCF-7 (low metastatic tumor), and MDA-MB-231 (metastatic tumor) cells were sampled at various time points (0 – 48 hrs) during EV harvest. These cell lines are widely studied to advance our understanding of the biology of breast cancer (BC).^52–54^ We found that, the number of exosomes, i.e. P_EV_, steadily increased with the harvest time but exhibited no significant difference among the cell lines (**Fig. 2d and Figure S9**). On the other hand, the P_miRNA_/P_EV_ ratio obtained by staining the EVs and labeling the enclosed miR-122 varied significantly between cell lines (**Fig. 2d**), with the EVs from MDA-MB-231 exhibiting the highest ratio and those from MCF-10A showing the lowest ratio. This result agrees with the previous findings on the specific loading of miR-122 in the exosomes from metastasis BC cell lines like MDA-MB-231.^55–57^

EVs present in human sera and their enclosed miRNAs were also analyzed by NOBEL-SPA. The samples were collected from BC patients and healthy controls. We captured the EVs via different surface proteins: the exosomal marker of CD63, and the tumor marker CD44, CD24, and HER2, to explore the loading of various tumor miRNAs (miR-21, miR-122, and miR-155) in these different EV sub-populations. A 24-microwell chip was fabricated on top of the cover glass to improve analysis throughput, with each well (max. volume = 10 µL) loaded with one type of the antibody-conjugated NOBs. Liquid mixing in each well was facilitated by the spinning function of the NOB; and the NOB can be immobilized by a magnet during washing and solution exchange. The highly abundant CD63^+^ exosomes required only 1 µL serum for detection, and 5 µL serum was used for capturing the EVs carrying each of the tumor proteins. A small set of clinical samples (n = 3 for each cohort of healthy controls or BC patients) was firstly analyzed (**Fig. 2e and Figure S10)**. We found that, neither the particle counts (P_EV_) nor the ratio of P_miRNA_/P_EV_ of the CD24^+^ EVs showed noticeable differences between BC patients and healthy controls (**Fig. S10a**). In contrast, a significantly higher P_EV_ was detected in BC patients for other EVs; and the CD44^+^ or HER2^+^ EVs showed larger differences between patients and healthy controls than the CD63^+^ EVs (**Fig. S10b-d**). Additionally, a much higher P_miRNA_/P_EV_ ratio of miR-122 or miR-21 was found in the CD63^+^, CD44^+^ or HER2^+^ EVs (**Fig. 2e**). But for miR-155, P_miRNA_ increased along with P_EV_, resulting in comparable P_miRNA_/P_EV_ ratio found in patients and controls. These results indicate that, secretion of miR-122- or miR-21 to the CD63^+^, CD44^+^ or HER2^+^ EVs was enhanced in BC patients, but the higher number of the miR-155-bearing EVs found in patients was just due to more total EVs present in the sera. Since a larger difference in the P_miRNA_/P_EV_ values between BC patients and healthy controls was found for miR-122 than for miR-21 within this small sample set, we analyzed the miR-122-associated CD63^+^ EVs in more clinical samples (n = 9 for each cohort). Indeed, not only P_EV_ and P_miR-122_ increased in the samples from BC patients (**Figure S11**), but also the P_miR-122_/P_EV_ ratio was significantly higher (*p* < 0.0001) in patients (**Fig. 2f**), agreeing with the previous reports on the higher amounts of exosomal miR-122 found in BC patients and the contribution of miR-122 to BC metastasis.^58, 59^ These results well justify the necessity of detecting the individual miRNA-associated EVs rather than the bulk EV quantity for recognition of unique EV sub-populations valuable for disease diagnosis.

### Assessment of colocalization of tumor proteins and miRNAs in EVs

The results discussed above point out that, miRNA loading in EVs carrying different surface markers could vary. Thus, it could be significant to detect both protein and miRNA simultaneously on the same EV. This is reasonable because the protein cargos are related to EV biogenesis pathways and reflective to the complex physiological states of the cells of origin,^21^ and the miRNAs could contribute to their roles in cell-cell communication.^16^ Dual-marker detection can be achieved by labeling each EV captured by the NOB with two fluorescent DNFs.^44, 45^ Five aptamer-containing ssDNA probes were designed to target the protein markers of CD63, CD44, HER2, EGFR, or MUC1. Each of these probes was paired with one of the four ssDNA probes containing the complementary sequence of miR-122, miR-21, let-7a, or miR-155 to assess their colocalization with the target proteins (**Figure S12a**). The upregulation of CD44, HER2, EGFR, and MUC1 has been widely reported in BC cells and tissues and in the EVs isolated from BC patients;^60–64^ and the miRNA targets also can regulate proliferation, migration, and invasion of breast cancer cells.^65, 66^ The ssDNA probes are RCA primers that can bind to the target and grow into DNF either labeled by Alexa 647 (for protein detection) or by Alexa 488 (for miRNA analysis). These two RCA systems exhibited very low crosstalk: mixing the mismatched primer and circular template did not initiate RCA (**Fig. S12b-k**).

**Figure 3a** displays the representative images obtained from dual detection of CD63 and miR-122 in the exosome standard (from COLO-1 cells) by NOBEL-SPA. We employed CellProfiler (https://cellprofiler.org/) for image analysis. This program recognizes the center of the fluorescence intensity of each fluorescent spot to determine the positive hits (i.e. one spot having only one exosome), counts the number of fluorescent spots in each channel, and measures their fluorescence intensities. Using the fluorescence intensity of Alexa633 (***I_protein_***) and Alexa488 (***I_miRNA_***), we can easily tell apart the exosomes carrying only the protein (red arrows in **Fig. 3a, *I_miRNA_*** = 0), only the miRNA (green arrows, ***I_protein_*** = 0), and both markers (yellow arrows). Only a small proportion of the CD63^+^ exosomes were associated with miR-122. Pairing CD63 with different miRNAs, colocalization analysis of the images gave out low coefficients ranging from 0.132 to 0.481, depending on the miRNA; and the P_miRNA_ counts obtained using the single-(for miRNA) and the dual-marker (for CD63 and miRNA) detection systems exhibited a high Pearson Correlation Coefficient of 0.9777 (**Figure S13**).

**Figure 3.**
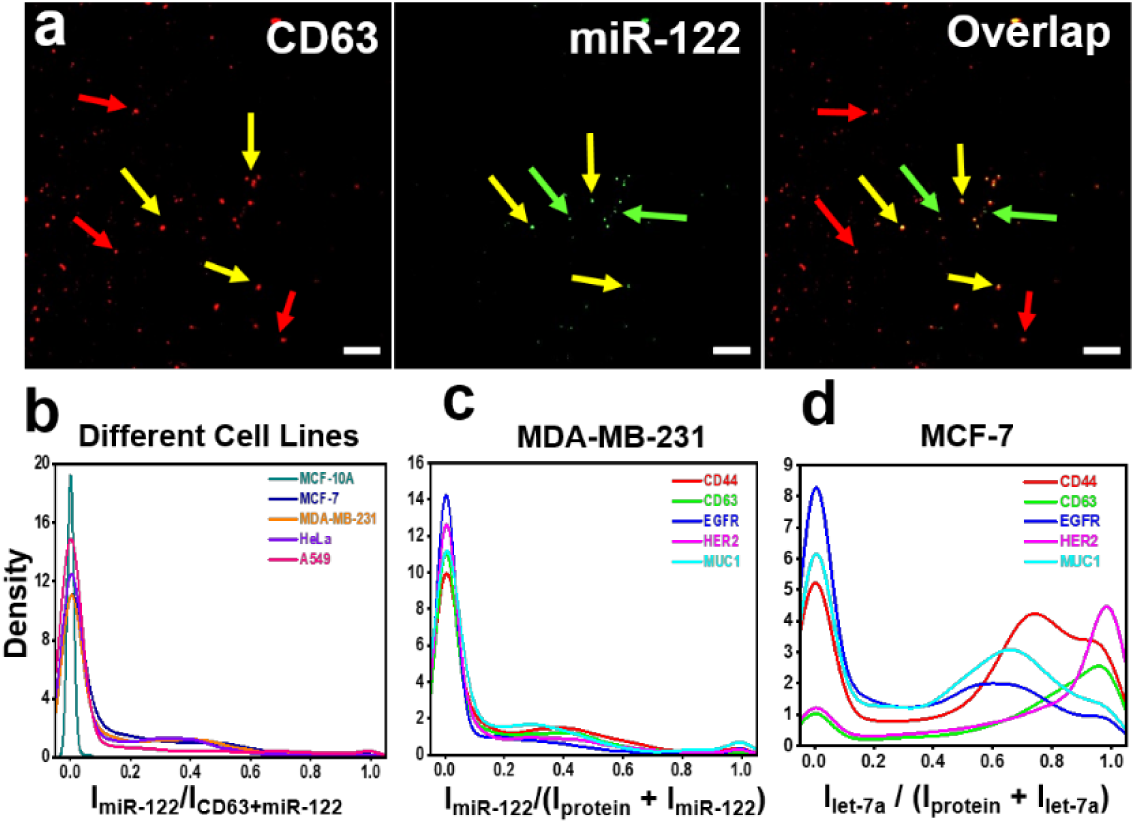
Dual-marker detection on single EVs. **a)** Representative CFM images for dual-marker detection of CD63 and miR-122 by NOBEL-SPA. Red, green, and yellow arrows point towards the particles emitting signals from only CD63, only miR-122, or both. **b-d)** Density distribution profiles of the fluorescence intensity ratios detected for the pair of CD63/miR-122 in 5 cell lines (**b**), for miR-122 in the exosomes from MDA-MB-231 (**c**), and for let-7a in those from MCF-7 with one of the 5 proteins (**d**).

The above results from standard exosomes prove the success of dual-labeling and low interference between the two DNF labels. Then we applied dual-label NOBEL-SPA to analyze the exosomes harvested from 5 cell lines: A549, HeLa, MCF-10A, MCF-7, and MDA-MB-231. A total of 20 protein/miRNA combinations were constituted from pairing 5 proteins and 4 miRNAs. In each EV sample from the same cell line, 2.4 × 10^6^ EVs were captured by ∼ 10^7^ NOB particles that target CD63/CD9/CD81; and 20 images were taken for each protein/miRNA pair. We plotted the density distribution profile of the ratio of ***I_miRNA_****/*(***I_protein_ + I_miRNA_***) for all the fluorescent spots detected in 20 images. Spots with a ratio of 0 or 1 were the EVs having only the protein or miRNA signal detected; and those with a ratio close to 0.5 should carry comparable amounts of protein and miRNA. We compared the distribution profiles of the same protein/miRNA pair among the EVs from different cell lines, or among the different protein/miRNA pairs from the EVs from the same cell line (**Figure S14-S18**). They well reflect the heterogenous nature of the EVs. Taking the CD63/miR-122 as one example (**Fig. 3b)**, we found that while miR-122 was not in the CD63**^+^** exosomes from MCF-10A cells, it was detected in the exosomes from MCF-7, MDA-MB-231 and HeLa cells. But even in these exosomes, miR-122 was only found in a very small proportion of the population. The colocalization situations of different protein/miRNA pairs in the exosomes derived from the same cell line were different as well. For example, slightly higher population densities were found for the MDA-MB-231 exosomes having miR-122 located with CD63, CD44, and MUC1 (i.e. the intensity ratios in the range of 0.4-0.6), compared to HER2 and EGFR (**Fig. 3c**); and a large proportion of the MCF-7 exosomes had let-7a with CD44, EGFR or MUC1, but not with HER2 or CD63 (**Fig. 3d**).

We segregated the exosomes into three categories: protein-only (*I_miRNA_ /(I_protein_ + I_miRNA_) = 0*), miRNA-only (*I_miRNA_ /(I_protein_ + I_miRNA_) = 1*), or dual-marker (0 < *I_miRNA_ /(I_protein_ + I_miRNA_) < 1*), counted the numbers of exosomes in each category, and calculated the proportion of each category among the total detected exosomes (sum of the particles found in all 3 categories). Fig. 4a is the heat map of the average (from 20 images) proportions of the dual-marker category detected in the EVs derived from 5 cell lines (5 columns), testing 20 protein/miRNA combinations (20 rows). Subjecting the proportions of the dual-marker category detected for each protein/miRNA pair (as one variable) in each image (as one repeat) to Canonical Discriminant Analysis (CDA), excellent classification of the exosomes by their cells of origin was achieved (Fig. 4b), with an error rate of 1% (Figure S19). The separation effect was worse if the proportions of the protein-only (Fig. 4c; and Fig. S20-21) or miRNA-only (Fig. S20 and S22) category was used in CDA.

**Figure 4.**
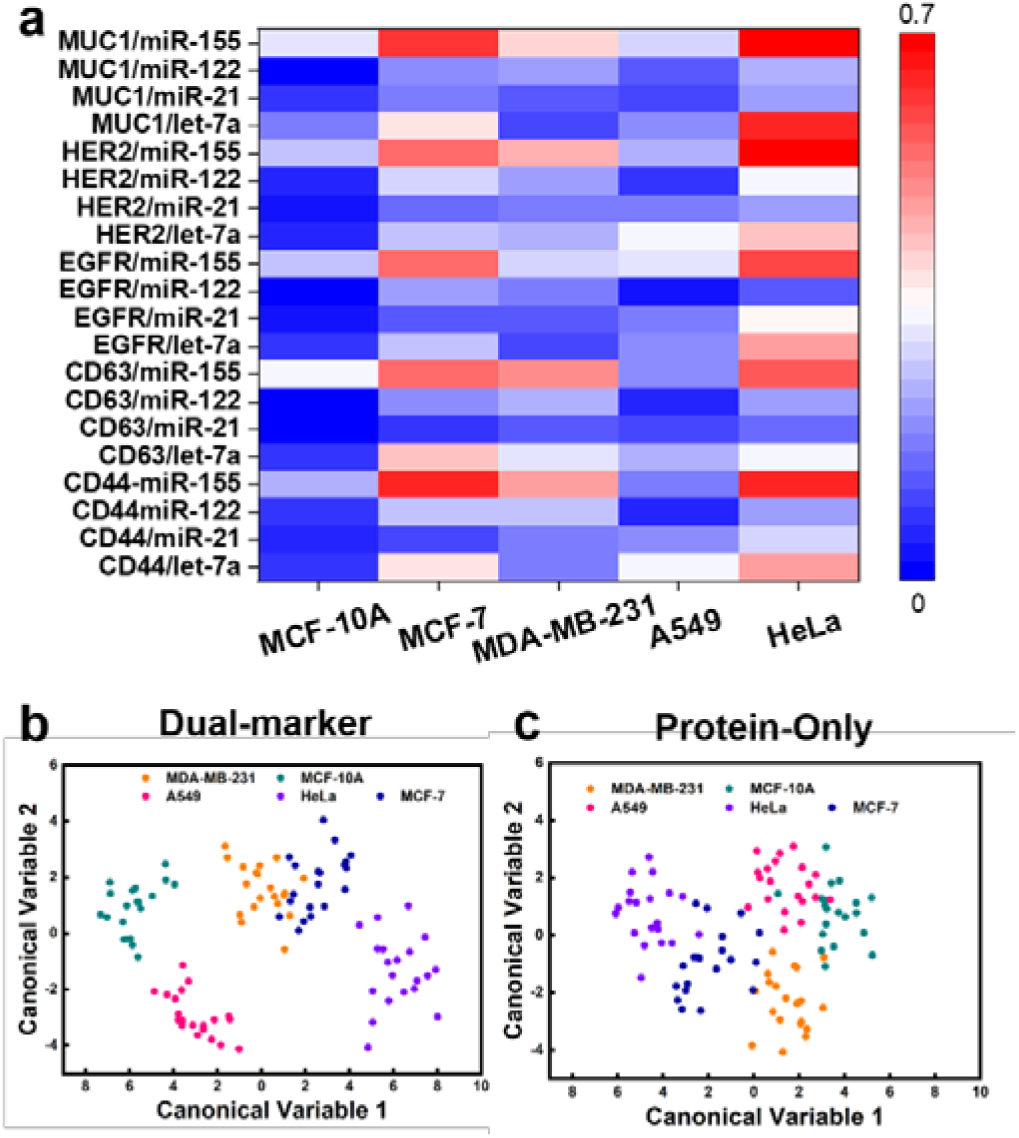
Differentiation of EVs by cells of origin based on the profiles of protein/miRNA combination. **a)** Heat map of the proportions of the protein^+^/miRNA^+^ exosomes among the total population detected in the EVs from different cells lines. **b-c)** Canonical Score Plots using the proportions of the dual-marker exosomes (**b**) or those with only proteins (**c**) showing different degrees of classification of the exosomes based on the cells of origin.

### EVs with dual markers in BC diagnosis

To reveal the most important protein/miRNA pairs for the differentiation effect, we analyzed the dataset with the machine learning algorithm of Support Vector Machine-Recursive Feature Elimination (**SVM-RFE**), as done in our other published works.^67, 68^ To find the protein/miRNA combinations suitable for BC diagnosis, we separated the five cell lines separated into four classes, non-BC tumors (A549 and HeLa), MCF-10A (non-tumor), MCF-7 (non-metastasis BC), and MDA-MB-231 (metastasis BC). Interestingly, 5 protein/miRNA pairs, CD44/miR-21, CD44/miR-155, CD63/miR-122, HER2/let-7a, and MUC1/let-7a, all belonging to the **dual-marker** category, were found to be the most important features for cell line differentiation. These 5 features are sufficient to classify the cell lines with satisfactory accuracy (0.800), specificity (0.927), sensitivity (0.800) and AUC (0.957), when tested by *10*-fold cross validation.

To further assess the power of the 5 protein/miRNA combinations found above, which represent 5 exosome sub-populations containing each specific marker pair, we employed NOBEL-SPA to examine human sera taken from Stage I (n = 11) and II (n = 9) BC patients, and healthy controls (n = 18) with matching ages. For each protein/miRNA pair, only ***1 µL human serum*** was used to mix with 10 µg NOBs conjugated with the anti-CD63/CD9/CD81 for exosome capture. Like in the cell line analysis, each CFM image was considered as one repeated measurement of the clinical sample; and each protein/miRNA pair considered as one variable in statistical analysis. From the violin plots of the proportion of each category: **protein-only**, **miRNA-only**, or **dual-marker** (**Fig. 5**), we found the proportions of both the CD63**^+^**/miR-122**^+^** and CD44**^+^**/miR-21**^+^** exosomes (enclosed in the two red rectangles in the dual-marker plot in **Fig. 5a**) increased significantly in BC patients compared to healthy controls. The CD63**^+^**/miR-122**^+^** sub-population even exhibited continuous and significant increase between healthy controls, Stage I, and Stage II patients (**Fig. 5c**). Such a gradual change was not observed for other sub-populations, although the CD44^+^ or HER2^+^ exosomes showed significant increase (p < 0.0001) in Stage I patients compared to healthy controls (**Fig. S23**). The good marker potential of these two proteins has also been revealed in our previous work.^45^ Interestingly, the abundance of the HER2^+^/let-7a^-^ exosomes increased in BC patients but that of the HER2^-^/let-7a^+^ dropped, which was also seen for the pair of CD44/miR-155 (enclosed in the blue rectangles in the protein-only and miRNA-only plot in **Fig. 5a**). The negative correlation could be related to the functions of these markers: HER2 can promote the growth of cancer cells, but let-7a can suppress migration and invasion of breast cancer cells.^69, 70^ Using the population proportions of the **protein^+^/miRNA^+^** exosomes observed in all images for *t*-distributed Stochastic Neighbor Embedding (*t*-SNE), the BC patients can be well differentiated from the health controls (**Fig. 5b**).

**Figure 5.**
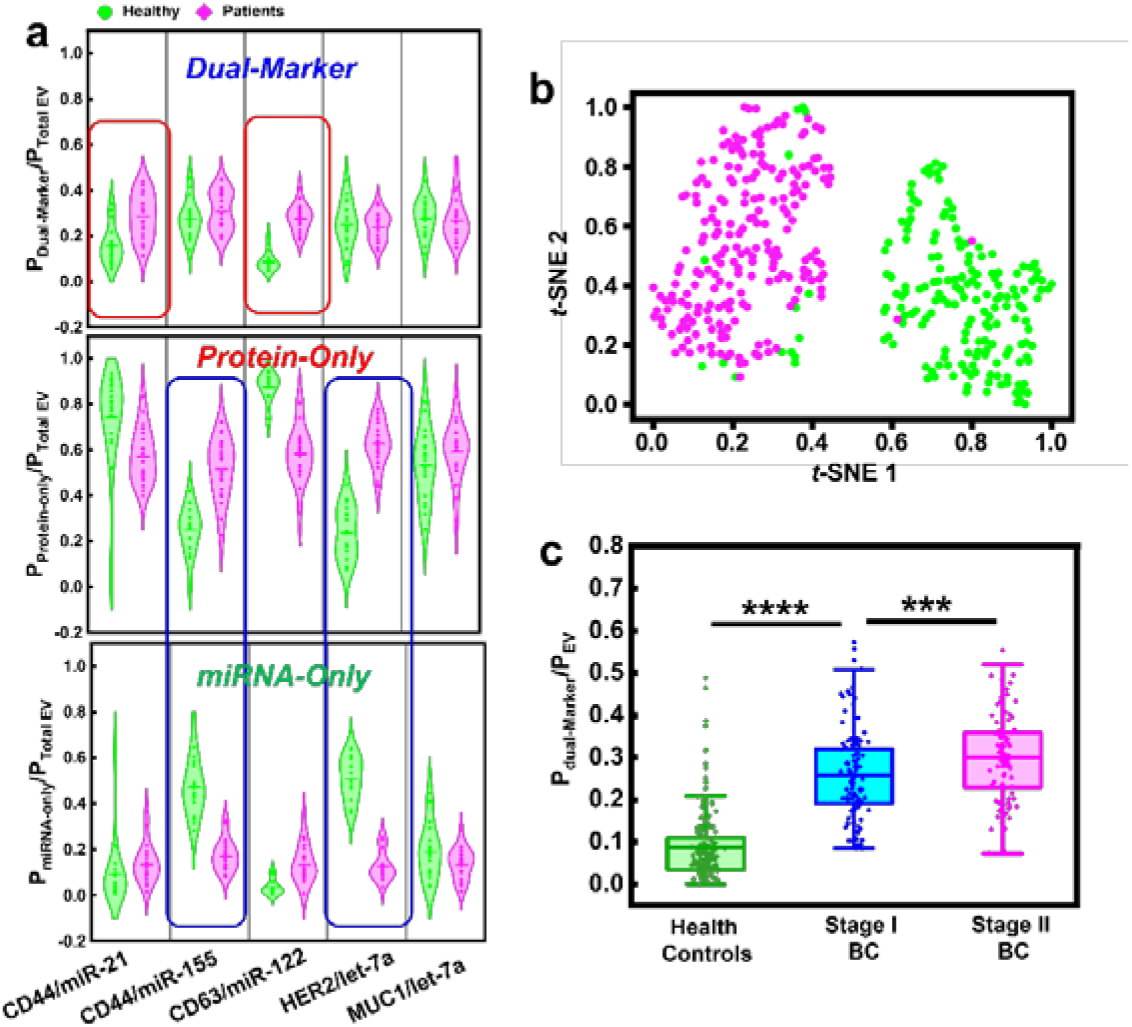
EV sub-populations for BC diagnosis. **a)** violin plots of the proportions of the different exosome sub-populations (protein^+^/miRNA^+^, protein^+^/miRNA^-^, and protein^-^/miRNA^+^) among the total exosomes detected in sera samples collected from healthy controls and BC patients. **b)** tSNE scatter plot showing successful differentiation of BC patients from healthy controls using the population proportions of the exosomes carrying dual markers of CD44/miR-21, CD44/miR-155, CD63/miR-122, HER2/let-7a, and MUC1/let-7a. Green dots represent healthy controls and the pink dots represent BC patients. **c)** Box plots of the proportions of the CD63^+^/miR-122^+^ exosome sub-populations among the total exosomes detected in sera samples collected from healthy controls, and Stage I or II BC patients.

## Conclusions

The present work has shown that NOBEL-SPA can realize easy and rapid single EV analysis and detect both the protein and RNA markers simultaneously. Its success relies on the unique structures of NOBs. The magnetic properties of NOBs permit easy handling and promote fast target binding and impurity removal. In addition, NOBs allow simple spatial separation of individual vesicle to reduce particle aggregation and enhance the confidence and easiness in single EV analysis. NOBEL-SPA can detect EVs in biological samples like cell culture medium and serum containing as low as 3-4 EV particles/µL in ∼ 4 hrs without any sample pre-processing. By immobilizing the intra-vesicular compounds on each NOB and visualizing single EVs, NOBEL-SPA can detect the low-abundance EV-enclosed miRNAs, even if fewer than one copy of the miRNA was found in each EV by RT-PCR; and can assess the colocalization of protein and miRNA markers in the same vesicle, which cannot be done by current techniques with matching assay efficiency and sensitivity (**Table S1**). More importantly, NOBEL-SPA can help unambiguous detection of the specific sub-populations from a swarm of heterogeneous EV in biofluids, which can enhance the accuracy in disease monitoring. Exploring the colocalization of protein and miRNA by NOBEL-SPA can also help reveal the loading mechanisms of miRNAs to EVs during disease development, leading to new therapeutic approaches by studying their production and functions. High heterogeneity ubiquitously exists in diverse biological particles including cells, bacteria, and viruses. While single particle analysis for those with dimensions larger than 1 micron can be easily done in instruments like flow cytometry, it remains challenging to study the cargo compositions in sub-micron particles like viruses and virus-like-particles. The working principle of NOBEL-SPA can be applied to the study of the sub-micron biological particles other than EVs to help gain more knowledge of their cargo loading and identify specific sub-populations defined by cargo combinations for diagnostic and therapeutic purposes.

## Experimental Section

### Preparation of bioconjugated NOBs

The magnetic Fe_3_O_4_@SiO_2_ nanorods were synthesized according to the reported protocol (Supporting Information).^71^ Then they were incubated with the carboxyl-modified (3-Aminopropyl)triethoxysilane (APTES) in dimethylformamide (DMF) at RT for 36 h, ready for antibody conjugation were done via EDC/NHS coupling. The remaining NHS ester on the surface will be deactivated by glycine. The obtained NOBs were redispersed in 1× PBS at a concentration of 1 mg/mL and stored at 4°C.

### NOBEL-SPA

The wells for the assay were firstly blocked by 0.1% BSA in 1× PBS overnight to reduce nonspecific adsorption. Ten µL of 10 ng/mL NOBs was added to each well and mixed with 10 µL of the EV sample. The chip was placed on the stirring plate set at 360 rpm. After 30-min EV capture, the NOBs were pulled down by a magnet and washed with 1× PBS, then sequentially mixed with 4% paraformaldehyde and 0.1 M EDC in 0.1 M imidazole buffer (pH 8) to fix the captured EV and crosslink the nucleic acid. The residual reagents were washed away with 0.2% glycine. Followed, a mixture of 1 µL 10× phi29, 1 µL of 0.125 mg/mL BSA, 1 µL of 0.05 µM circular probe, 1 µL of 0.25 µM recognition probe, 1 µL 0.5x DiB, and 6 µL DI water were added and incubated for 30 min at room temperature on the magnetic stir plate (360 rpm) to recognize the target miRNA and stain the captured EVs. After that, 1 µL of 200 µM dNTP, 1 µL of 2.5 µM biotin-dATP, and 1 µL phi29 DNA polymerase (2.5 Units/mL) were added to the well and incubated at 37 °C for 30min. At last, 1 µL of 2.5 µM streptavidin-modified Alexa 633 was added and the solution was incubated for 30 min. After washed with 1× PBS three times and dispersed in 10 µL 1× PBS, the NOBs were ready for CFM. Dual-marker NOBEL-SPA followed a similar procedure, the details of which can be found in Supporting Information.

### Confocal microscopy and image analysis

Fluorescence imaging was performed on a Zeiss 880 Inverted Confocal Microscope using a UV laser with λ_ex_ = 330 nm, an Argon laser with λ_ex_ = 488 nm, and a HeNe laser at λ_ex_ = 633 nm for fluorescence from DiB, Alexa 488, and Alexa 633 (or Alexa 647), respectively. All CFM images were collected at a resolution of 512 × 512 pixels. The viewing area was 100 μm × 100 μm. For each sample, 10 images were acquired at 10 different locations from a single well. The raw data obtained from CFM were exported to the tiff format via ZEN 3.2 (blue edition). The images were processed by CellProfiler with a lab-built pipeline. In this pipeline, the clumped cluster was divided by the intensity and the size smaller than 20 pixel was regarded as one spot. The number of particles, the size and fluorescence intensity detected on each particle were automatically collected by the pipeline. More details can be found in Supporting Information.

### Statistical analysis

Data plots, statistical analysis, and canonical classification were carried out by Origin 2021. Differences with *p* < 0.05 were considered as statistically significant. The *t*-Distributed Stochastic Neighbor Embedding (*t*-SNE) was performed to reduce the dimensionality of complex data by Python 3.9 (64-bit) with the following parameters: n_components=2, init=’pca’, verbose=1, random_state=123, perplexity=15, learning_rate=’auto’, n_iter=5000. The two dimension data was plotted by the matplotlib.pyplot. Feature selection and classification were performed with Python 3.9 (64-bit), using StandardScaler for data standardization, Support Vector Machine-Recursive Feature Elimination (SVM-RFE) to select the top 5 sensors according to weight vectors by the iteration process of the backward removal of features, RFE (estimator = svm.SVC (kernel = &#39; linear&#39), n_features_to_select=5). Performance metrics for the classification evaluation were calculated by using RepeatedStratifiedKFold (n_splits = 10, n_repeats = 3) for cross validation and with svm.SVC (kernel=&#39; linear&#39) as the estimator. All data points derived from each experiment was collected with multiple repeats.

## ASSOCIATED CONTENT

### Author Information

#### Authors

**Zongbo Li –** Department of Chemistry, University of California-Riverside, CA 92521, USA.

**Kaizhu Guo –** Department of Chemistry, University of California-Riverside, CA 92521, USA.

**Ziting Gao –** Department of Chemistry, University of California-Riverside, CA 92521, USA.

**Junyi Chen –** Environmental Toxicology Graduate Program, University of California-Riverside, CA 92521, USA.

**Zuyang Ye –** Department of Chemistry, University of California-Riverside, CA 92521, USA.

**Shizhen Emily Wang** – Professor of Pathology, University of California-San Diego, La Jolla, CA 92093, USA <u>orcid.org/0000-0003-0218-3042</u>

**Yadong Yin** – Professor of Chemistry, University of California-Riverside, Riverside, CA 92521, USA. <u>orcid.org/0000-0002-3317-3464</u>

### Author Contributions

W.Z. conceived the original idea. Z.L. and K.G. performed most of the experiments and data analysis. Z.G. prepared the cells and EVs. J.C. assisted with the statistical analysis. Z.Y. and Y.Y. helped with material synthesis. S.E.W. provided guidance on cell line and marker selections. W.Z. and Z.L. wrote the entire manuscript. Z.L. wrote the experimental details and results. All authors have given approval to the final version of the manuscript.

## Acknowledgements

The authors would like to thank UCR’s Extramural Funding Opportunity Preparation Award (EFOPA), and City of Hope – UC Riverside Biomedical Research Initiative (CUBRI) Award to W. Zhong. S. E. Wang acknowledges the support by the National Institutes of Health (NIH)/National Cancer Institute (NCI) under grant number R01CA266486. Y. Yin and Z. Ye were supported by the Engineering Research Centers Program of the National Science Foundation under NSF Cooperative Agreement No. 1941543. Electron microscopy was performed on a scanning electron microscope NNS450 instrument in the Central Facility for Advanced Microscopy and Microanalysis (CFAMM) at UC Riverside, with the assistance of Dr. ILkeun Lee.

## Supporting Information Available

Additional experimental details; supplementary table for summary of current methods; additional material characterization and methodology validation results; information of the sequences designed and employed; data for specificity test; additional data, including images and density distribution plots for analysis of EVs derived from cells, and the corresponding heatmaps and classification results; and additional data and statistical analysis results and plots for EVs in human sera.

## Table of Contents

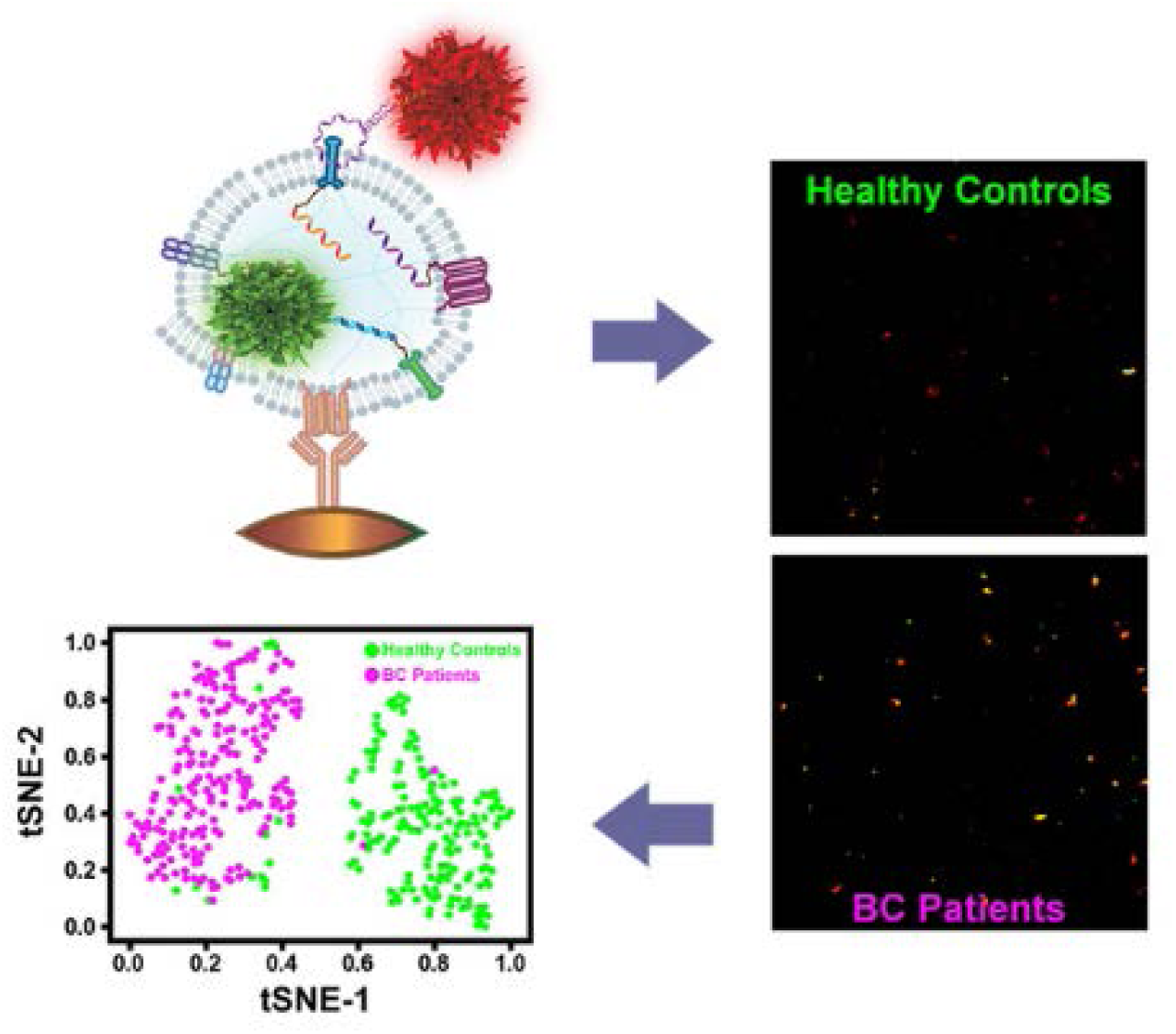

## Notes

### Competing Interest Statement

The authors have declared no competing interest.

## References

(1) Iraci, N.; Leonardi, T.; Gessler, F.; Vega, B.; Pluchino, S. Focus on extracellular vesicles: physiological role and signalling properties of extracellular membrane vesicles. Int. J. Mol. Sci. 2016, *17*, 171/171–171/132.

(2) Kraemer-Albers, E.-M.; Hill, A. F. Extracellular vesicles: interneural shuttles of complex messages. Curr. Opin. Neurobiol. 2016, 39, 101–107.

(3) Shifrin, D. A., Jr.; Beckler, M. D.; Coffey, R. J.; Tyska, M. J. Extracellular vesicles: communication, coercion, and conditioning. Mol. Biol. Cell 2013, 24, 1253–1259.

(4) Turturici, G.; Tinnirello, R.; Sconzo, G.; Geraci, F. Extracellular membrane vesicles as a mechanism of cell-to-cell communication: advantages and disadvantages. Am. J. Physiol. 2014, 306, C621–C633.

(5) Yanez-Mo, M.; Siljander, P. R. M.; Andreu, Z.; Zavec, A. B.; Borras, F. E.; Buzas, E. I.; Buzas, K.; Casal, E.; Cappello, F.; Carvalho, J.;, et al. Biological properties of extracellular vesicles and their physiological functions. J. Extracell. Vesicles 2015, 4, 27066–27066.

(6) Crowley, E.; Di Nicolantonio, F.; Loupakis, F.; Bardelli, A. Liquid biopsy: monitoring cancer-genetics in the blood. Nat. Rev. Clin. Oncol. 2013, 10, 472–484.

(7) Lo Cicero, A.; Stahl, P. D.; Raposo, G. Extracellular vesicles shuffling intercellular messages: for good or for bad. Curr. Opin. Cell Biol. 2015, 35, 69–77.

(8) Hoshino, A.; Kim, H. S.; Bojmar, L.; Gyan, K. E.; Cioffi, M.; Hernandez, J.; Zambirinis, C. P.; Rodrigues, G.; Molina, H.; Heissel, S.;, et al. Extracellular Vesicle and Particle Biomarkers Define Multiple Human Cancers. Cell 2020, 182, 1044–1061.e1018.

(9) Cocucci, E.; Meldolesi, J. Ectosomes. Curr. Biol. 2011, 21, R940–R941.

(10) Cocucci, E.; Racchetti, G.; Meldolesi, J. Shedding microvesicles: artefacts no more. Trends Cell Biol. 2009, 19, 43–51.

(11) Hanson, P. I.; Cashikar, A. Multivesicular body morphogenesis. Annu. Rev. Cell Dev. Biol. 2012, 28, 337–362.

(12) Quesenberry, P. J.; Goldberg, L. R.; Aliotta, J. M.; Dooner, M. S.; Pereira, M. G.; Wen, S.; Camussi, G. Cellular phenotype and extracellular vesicles: basic and clinical considerations. Stem Cells and Dev. 2014, 23, 1429–1436.

(13) Raposo, G.; Stoorvogel, W. Extracellular vesicles: exosomes, microvesicles, and friends. J. Cell Biol. 2013, 200, 373–383.

(14) Teis, D.; Saksena, S.; Emr, S. D. SnapShot: the ESCRT machinery. Cell 2009, 137, 182–182.e181.

(15) Tian, T.; Wang, Y.; Wang, H.; Zhu, Z.; Xiao, Z. Visualizing of the cellular uptake and intracellular trafficking of exosomes by live-cell microscopy. J. Cell Biochem. 2010, 111, 488–496.

(16) Pluchino, S.; Smith, J. A. Explicating Exosomes: Reclassifying the Rising Stars of Intercellular Communication. Cell 2019, 177, 225–227.

(17) Atayde, V. D.; Hassani, K.; da Silva Lira Filho, A.; Borges, A. R.; Adhikari, A.; Martel, C.; Olivier, M. Leishmania exosomes and other virulence factors: Impact on innate immune response and macrophage functions. Cellul. immunol. 2016, 309, 7–18.

(18) Broekman, M. L.; Maas, S. L. N.; Abels, E. R.; Mempel, T. R.; Krichevsky, A. M.; Breakefield, X. O. Multidimensional communication in the microenvirons of glioblastoma. Nature reviews. Neurol. 2018, 14, 482–495.

(19) Wang, Y.-M.; Trinh, M. P.; Zheng, Y.; Guo, K.; Jimenez, L. A.; Zhong, W. Analysis of circulating non-coding RNAs in a non-invasive and cost-effective manner. Trends Anal. Chem. 2019, 117, 242–262.

(20) Schorey, J. S.; Cheng, Y.; Singh, P. P.; Smith, V. L. Exosomes and other extracellular vesicles in host-pathogen interactions. EMBO reports 2015, 16, 24–43.

(21) Kalluri, R.; LeBleu, V. S. The biology, function, and biomedical applications of exosomes. Science 2020, 367, eaau6977.

(22) van Niel, G.; D’Angelo, G.; Raposo, G. Shedding light on the cell biology of extracellular vesicles. Nat. Rev. Mol. Cell Biol. 2018, 19, 213–228.

(23) Maas, S. L. N.; Breakefield, X. O.; Weaver, A. M. Extracellular Vesicles: Unique Intercellular Delivery Vehicles. Trends Cell Biol. 2017, 27, 172–188.

(24) Wang, S.; Khan, A.; Huang, R.; Ye, S.; Di, K.; Xiong, T.; Li, Z. Recent advances in single extracellular vesicle detection methods. Biosen. Bioelectron. 2020, 154, 112056.

(25) Cocozza, F.; Grisard, E.; Martin-Jaular, L.; Mathieu, M.; Théry, C. SnapShot: Extracellular Vesicles. Cell 2020, 182, 262–262.e261.

(26) Zhang, P.; Zhou, X.; Zeng, Y. Multiplexed immunophenotyping of circulating exosomes on nano-engineered ExoProfile chip towards early diagnosis of cancer. Chem. Sci. 2019, 10, 5495–5504.

(27) Ferguson, S.; Weissleder, R. Modeling EV Kinetics for Use in Early Cancer Detection. Adv. Biosys. 2020, 4, 1900305.

(28) Morales-Kastresana, A.; Musich, T. A.; Welsh, J. A.; Telford, W.; Demberg, T.; Wood, J. C. S.; Bigos, M.; Ross, C. D.; Kachynski, A.; Dean, A.;, et al. High-fidelity detection and sorting of nanoscale vesicles in viral disease and cancer. J. Extracell. Vesicles 2019, 8, 1597603.

(29) Panagopoulou, M. S.; Wark, A. W.; Birch, D. J. S.; Gregory, C. D. Phenotypic analysis of extracellular vesicles: a review on the applications of fluorescence. J. Extracell. Vesicles 2020, 9, 1710020.

(30) Tian, Y.; Gong, M.; Hu, Y.; Liu, H.; Zhang, W.; Zhang, M.; Hu, X.; Aubert, D.; Zhu, S.; Wu, L.;, et al. Quality and efficiency assessment of six extracellular vesicle isolation methods by nano-flow cytometry. J. Extracell. Vesicles 2020, 9, 1697028.

(31) Welsh, J. A.; Van Der Pol, E.; Arkesteijn, G. J. A.; Bremer, M.; Brisson, A.; Coumans, F.; Dignat-George, F.; Duggan, E.; Ghiran, I.; Giebel, B.;, et al. MIFlowCyt-EV: a framework for standardized reporting of extracellular vesicle flow cytometry experiments. J. Extracell. Vesicles 2020, 9, 1713526.

(32) Nizamudeen, Z.; Markus, R.; Lodge, R.; Parmenter, C.; Platt, M.; Chakrabarti, L.; Sottile, V. Rapid and accurate analysis of stem cell-derived extracellular vesicles with super resolution microscopy and live imaging. Biochim Biophys Acta Mol Cell Res 2018, 1865, 1891–1900.

(33) McNamara, R. P.; Zhou, Y.; Eason, A. B.; Landis, J. T.; Chambers, M. G.; Willcox, S.; Peterson, T. A.; Schouest, B.; Maness, N. J.; MacLean, A. G.;, et al. Imaging of surface microdomains on individual extracellular vesicles in 3-D. J. Extracell. Vesicles 2022, 11, e12191.

(34) Wu, D.; Yan, J.; Shen, X.; Sun, Y.; Thulin, M.; Cai, Y.; Wik, L.; Shen, Q.; Oelrich, J.; Qian, X.;, et al. Profiling surface proteins on individual exosomes using a proximity barcoding assay. Nat. Comm. 2019, 10, 3854.

(35) Lee, K.; Fraser, K.; Ghaddar, B.; Yang, K.; Kim, E.; Balaj, L.; Chiocca, E. A.; Breakefield, X. O.; Lee, H.; Weissleder, R. Multiplexed Profiling of Single Extracellular Vesicles. ACS Nano 2018, 12, 494–503.

(36) Min, J.; Son, T.; Hong, J.-S.; Cheah, P. S.; Wegemann, A.; Murlidharan, K.; Weissleder, R.; Lee, H.; Im, H. Plasmon-Enhanced Biosensing for Multiplexed Profiling of Extracellular Vesicles. Adv. Biosys. 2020, 2000003.

(37) Jeong, M. H.; Son, T.; Tae, Y. K.; Park, C. H.; Lee, H. S.; Chung, M. J.; Park, J. Y.; Castro, C. M.; Weissleder, R.; Jo, J. H.;, et al. Plasmon-Enhanced Single Extracellular Vesicle Analysis for Cholangiocarcinoma Diagnosis. Adv. Sci. 2023, e2205148.

(38) Wang, S.; Zheng, W.; Wang, R.; Zhang, L.; Yang, L.; Wang, T.; Saliba, J. G.; Chandra, S.; Li, C.-Z.; Lyon, C. J.;, et al. Monocrystalline Labeling Enables Stable Plasmonic Enhancement for Isolation-Free Extracellular Vesicle Analysis. Small 2023, 19, 2204298.

(39) Wei, P.; Wu, F.; Kang, B.; Sun, X.; Heskia, F.; Pachot, A.; Liang, J.; Li, D. Plasma extracellular vesicles detected by Single Molecule array technology as a liquid biopsy for colorectal cancer. J Extracell Vesicles 2020, 9, 1809765.

(40) He, D.; Wang, H.; Ho, S. L.; Chan, H. N.; Hai, L.; He, X.; Wang, K.; Li, H. W. Total internal reflection-based single-vesicle in situ quantitative and stoichiometric analysis of tumor-derived exosomal microRNAs for diagnosis and treatment monitoring. Theranostics 2019, 9, 4494–4507.

(41) Zhou, J.; Wu, Z.; Hu, J.; Yang, D.; Chen, X.; Wang, Q.; Liu, J.; Dou, M.; Peng, W.; Wu, Y.;, et al. High-throughput single-EV liquid biopsy: Rapid, simultaneous, and multiplexed detection of nucleic acids, proteins, and their combinations. Sci. Adv. 2020, 6.

(42) Ayers, L.; Pink, R.; Carter, D. R. F.; Nieuwland, R. Clinical requirements for extracellular vesicle assays. J. Extracell. Vesicles 2019, 8, 1593755.

(43) Guo, K. Z.; Li, Z. B.; Win, A.; Coreas, R.; Adkins, G. B.; Cui, X. P.; Yan, D.; Cao, M. H.; Wang, S. E.; Zhong, W. W. Calibration-free analysis of surface proteins on single extracellular vesicles enabled by DNA nanostructure. Biosens Bioelectron 2021, 192.

(44) Shen, W.; Guo, K.; Adkins, G. B.; Jiang, Q.; Liu, Y.; Sedano, S.; Duan, Y.; Yan, W.; Wang, S. E.; Bergersen, K.;, et al. A Single Extracellular Vesicle (EV) Flow Cytometry Approach to Reveal EV Heterogeneity. Angew. Chem. Int. Ed. 2018, 57, 15675–15680.

(45) Guo, K.; Li, Z.; Win, A.; Coreas, R.; Adkins, G. B.; Cui, X.; Yan, D.; Cao, M.; Wang, S. E.; Zhong, W. Calibration-free analysis of surface proteins on single extracellular vesicles enabled by DNA nanostructure. Biosen. Bioelectron. 2021, 192, 113502.

(46) Wang, M.; He, L.; Xu, W.; Wang, X.; Yin, Y. Magnetic assembly and field-tuning of ellipsoidal-nanoparticle-based colloidal photonic crystals. Angew. Chem. Int. ed. 2015, 54, 7077–7081.

(47) Li, B.; Chen, J.; Han, L.; Bai, Y.; Fan, Q.; Wu, C.; Wang, X.; Lee, M.; Xin, H. L.; Han, Z.;, et al. Ligand-Assisted Solid-State Transformation of Nanoparticles. Chem. Mat. 2020, 32, 3271–3277.

(48) Xu, W.; Wang, M.; Li, Z.; Wang, X.; Wang, Y.; Xing, M.; Yin, Y. Chemical Transformation of Colloidal Nanostructures with Morphological Preservation by Surface-Protection with Capping Ligands. Nano Letters 2017, 17, 2713–2718.

(49) Gupta, M. P.; Tandalam, S.; Ostrager, S.; Lever, A. S.; Fung, A. R.; Hurley, D. D.; Alegre, G. B.; Espinal, J. E.; Remmel, H. L.; Mukherjee, S.;, et al. Non-reversible tissue fixation retains extracellular vesicles for in situ imaging. Nat. Met. 2019, 16, 1269–1273.

(50) Pena, J. T. G.; Sohn-Lee, C.; Rouhanifard, S. H.; Ludwig, J.; Hafner, M.; Mihailovic, A.; Lim, C.; Holoch, D.; Berninger, P.; Zavolan, M.;, et al. miRNA in situ hybridization in formaldehyde and EDC–fixed tissues. Nat. Met. 2009, 6, 139–141.

(51) Cui, L.; Peng, R. X.; Zeng, C. F.; Zhang, J. L.; Lu, Y. Z.; Zhu, L.; Huang, M. J.; Tian, Q. H.; Song, Y. L.; Yang, C. Y. A general strategy for detection of tumor-derived extracellular vesicle microRNAs using aptamer-mediated vesicle fusion. Nano Today 2022, 46.

(52) Holliday, D. L.; Speirs, V. Choosing the right cell line for breast cancer research. Breast Cancer Res. 2011, 13, 211–217.

(53) Neve, R. M.; Chin, K.; Fridlyand, J.; Yeh, J.; Baehner, F. L.; Fevr, T.; et al. A collection of breast cancer cell lines for the study of functionally distinct cancer subtypes. Cancer Cell 2006, 10, 515–527.

(54) Dai, X.; Cheng, H.; Bai, Z.; Li, J. Breast Cancer Cell Line Classification and Its Relevance with Breast Tumor Subtyping. J. Cancer 2017, 18, 3131–3141.

(55) Cao, M.; Isaac, R.; Yan, W.; Ruan, X.; Jiang, L.; Wan, Y.; Wang, J.; Wang, E.; Caron, C.; Neben, S.;, et al. Cancer-cell-secreted extracellular vesicles suppress insulin secretion through miR-122 to impair systemic glucose homeostasis and contribute to tumour growth. Nat. Cell Biol. 2022, 24, 954–967.

(56) Fong, M. Y.; Zhou, W.; Liu, L.; Alontaga, A. Y.; Chandra, M.; Ashby, J.; Chow, A.; O’Connor, S. T. F.; Li, S.; Chin, A. R.;, et al. Breast-cancer-secreted miR-122 reprograms glucose metabolism in premetastatic niche to promote metastasis. Nat. Cell Biol. 2015, 17, 183–194.

(57) Temoche-Diaz, M. M.; Shurtleff, M. J.; Nottingham, R. M.; Yao, J.; Fadadu, R. P.; Lambowitz, A. M.; Schekman, R. Distinct mechanisms of microRNA sorting into cancer cell-derived extracellular vesicle subtypes. eLife 2019, 8, e47544.

(58) Fong, M. Y.; Zhou, W. Y.; Liu, L.; Alontaga, A. Y.; Chandra, M.; Ashby, J.; Chow, A.; O’Connor, S. T. F.; Li, S. S.; Chin, A. R.;, et al. Breast-cancer-secreted miR-122 reprograms glucose metabolism in premetastatic niche to promote metastasis. Nat Cell Biol 2015, 17, 183-+.

(59) Cao, M. H.; Isaac, R.; Yan, W.; Ruan, X. H.; Jiang, L.; Wan, Y. H.; Wang, J.; Wang, E.; Caron, C.; Neben, S.;, et al. Cancer-cell-secreted extracellular vesicles suppress insulin secretion through miR-122 to impair systemic glucose homeostasis and contribute to tumour growth. Nat Cell Biol 2022, 24, 954-+.

(60) Jerabkova-Roda, K.; Dupas, A.; Osmani, N.; Hyenne, V.; Goetz, J. G. Circulating extracellular vesicles and tumor cells: sticky partners in metastasis. Trends Cancer 2022, 8, 799–805.

(61) Ekstrom, K.; Crescitelli, R.; Petursson, H. I.; Johansson, J.; Lasser, C.; Olofsson Bagge, R. Characterization of surface markers on extracellular vesicles isolated from lymphatic exudate from patients with breast cancer. BMC Cancer 2022, 22, 50.

(62) Tian, F.; Zhang, S.; Liu, C.; Han, Z.; Liu, Y.; Deng, J.; Li, Y.; Wu, X.; Cai, L.; Qin, L.;, et al. Protein analysis of extracellular vesicles to monitor and predict therapeutic response in metastatic breast cancer. Nat Commun 2021, 12, 2536.

(63) Hollingsworth, M. A.; Swanson, B. J. Mucins in cancer: protection and control of the cell surface. Nat Rev Cancer 2004, 4, 45–60.

(64) Li, R.; An, Y.; Jin, T. Y.; Zhang, F.; He, P. G. Detection of MUC1 protein on tumor cells and their derived exosomes for breast cancer surveillance with an electrochemiluminescence aptasensor. J Electroanal Chem 2021, 882.

(65) Abolghasemi, M.; Tehrani, S. S.; Yousefi, T.; Karimian, A.; Mahmoodpoor, A.; Ghamari, A.; Jadidi-Niaragh, F.; Yousefi, M.; Kafil, H. S.; Bastami, M.;, et al. MicroRNAs in breast cancer: Roles, functions, and mechanism of actions. J Cell Physiol 2020, 235, 5008–5029.

(66) Li, M. H.; Zou, X.; Xia, T. S.; Wang, T. S.; Liu, P.; Zhou, X.; Wang, S.; Zhu, W. A five-miRNA panel in plasma was identified for breast cancer diagnosis. Cancer Med-Us 2019, 8, 7006–7017.

(67) Chen, J.; Gill, A. D.; Hickey, B. L.; Gao, Z.; Cui, X.; Hooley, R. J.; Zhong, W. Machine Learning Aids Classification and Discrimination of Noncanonical DNA Folding Motifs by an Arrayed Host:Guest Sensing System. J. Am. Chem. Soc. 2021, 143, 12791–12799.

(68) Chen, J.; Hickey, B. L.; Gao, Z.; Raz, A. A. P.; Hooley, R. J.; Zhong, W. Sensing Base Modifications in Non-Canonically Folded DNA with an Optimized Host:Guest Sensing Array. ACS sensors 2022, 7, 2164–2169.

(69) Yarden, Y. Biology of HER2 and its importance in breast cancer. Oncology-Basel 2001, 61, 1–13.

(70) Kim, S. J.; Shin, J. Y.; Lee, K. D.; Bae, Y. K.; Sung, K. W.; Nam, S. J.; Chun, K. H. MicroRNA let-7a suppresses breast cancer cell migration and invasion through downregulation of C-C chemokine receptor type 7. Breast Cancer Res 2012, 14.

(71) Li, Z. W.; Jin, J. B.; Yang, F.; Song, N. N.; Yin, Y. D. Coupling magnetic and plasmonic anisotropy in hybrid nanorods for mechanochromic responses. Nat. Comm. 2020, 11.

