## supplemental file for "Colocalization of Protein and microRNA Markers Reveals Unique Extracellular Vesicle Sub-Populations for Early Cancer Detection"

†: Authors contributed equally to the work.

### Supporting Information

#### Table of Contents

**Chemicals and Reagents** ........ **S3**

**Supplementary Methods** ........ **S4**

**Supplementary Table** ........ **S9**

**Supplementary Figures** ........ **S10**

#### Chemicals and Reagents

Iron(III) chloride hexahydrate (FeCl3.6H2O, 99+%, extra pure, A0399225), Tetraethyl orthosilicate(TEOS, 42036100), succinic anhydride(AC15876-0500), ethanolamine (427251000), paraformaldehyde, 96% (AC41678-5000) were purchased from Acros Organics. Poly(acrylic acid) (PAA ~1800, 323667-100G), (3-Aminopropyl)triethoxysilane (APTES, 99%, 440140-100ML), albumin (A9511-500MG, ≥97%), glycine (G8898-500G, ≥99%), triethylene glycol (TEG, T59455-1KG) anti-CD9 antibody (CBL162), and anti-CD81 antibody (SAB4700232-100UG) were purchased from Sigma Aldrich. Ammonium solution, 1-ethyl-3-(-3-dimethylaminopropyl) carbodiimide hydrochloride (EDC, 22980), N-hydroxysuccinimide (NHS, 24500), goat anti-mouse IgG (H+L,horseradish peroxidase conjugate, cross-adsorbed) (A16072) were purchased from the ThermoFisher Scientific. N,N-dimethylformamide (DX1730-6, DMF) was purchased from the EMD. Sodium borate (S249-500), sodium chloride (BP358-1, NaCl), MES (BP300-100), Tris base (BP152-500), sulfuric acid, 98% (A300S-500), and hydrogen peroxide, 30% (H325-500), phosphate buffered saline (PBS), 10× solution (AAJ75889K8, molecular biology grade; consisting of 80.6 mM sodium phosphate, 19.4 mM potassium phosphate, 27 mM KCl, and 1.37 M NaCl in high-purity deionized H2O), and anti-HER2 antibody (MAB1129100) were purchased from the Fisher Scientific. The phi29 DNA Polymerase (M0296L), T4 DNA ligase (M020S), exonuclease I (M0293S), exonuclease III (M0206S) were purchased from the New England Biolabs (NEB). Glutaraldehyde(A17876) was purchased from Alfa Aesar. Anti-CD63 antibody (NB100-77913), and anti-CD44 antibody (NBP2-46151) was purchased from the Novus Biologicals. Anti-CD24 antibody (311101), biotinylated mouse antihuman CD63 antibody (353017) were purchased from the BioLegend. The streptavidin horseradish peroxidase (HRP) (10755) was purchased from the Cepham Life Sciences. The SYLGARD™ 184 Silicone Elastomer Kit was purchased from the Dow Inc. Purified EVs (lyophilized exosomes from human colon carcinoma cell line COLO-1) were purchased from HansaBioMed Life Science Ltd. (Tallinn, Estonia). The CellBrite Blue Cytoplasmic Membrane-Labeling Kit (DiB, 30024) was purchased from Biotium. Human breast cancer patient serum samples and healthy patient serum samples were provided by the NCI funded Cooperative Human Tissue Network (CHTN).

### Supplementary Methods

#### 1. Cell culture and EV isolation

All cells were cultured at the recommended media containing 1% penicillin streptomycin. MCF-10A cells were cultured in the DMEM/F-12 media supplied with 5% horse serum, 0.1 μg/mL cholera toxin, 10 μg/mL insulin, 0.5 μg/mL hydrocortisone, and 20 ng/mL EGF. HeLa, A549, MCF-7 and MDA-MB-231 cells were cultured in DMEM media supplemented with 10% FBS. All cell lines were maintained at 37 °C in a humidified 5% $CO_2$ incubator and routinely screened for Mycoplasa contamination. When the cells reached a confluency of 75%, the medium was replaced with the EV-depleted culture medium. After 24 h incubation, the culture medium was collected and centrifuged at 500 *g* for 15 min and 15,000 *g* for 20 min to remove the cell debris. Next, the medium was ultra-centrifuged at 110,000 *g* for 19 h to pellet the EVs secreted by the cells. The EVs pellet was resuspended in 1 × PBS. Particle concentration in the EV solution was measured by NTA.

#### 2. Synthesis of silica-coated magnetic nanorods ($Fe_3O_4$@$SiO_2$)

First, 10.8 g of $FeCl_3 \cdot 6H_2O$ was dissolved in 400 mL deionized water (DI water) and incubated at 87 °C for 18 h. The precipitation was collected and washed with DI water three times at 11,000 rpm for 15 minutes. The obtained FeOOH rods were dispersed in 40 mL DI water. To coat the FeOOH with $SiO_2$, the FeOOH rods were modified with PAA firstly via incubating 10 mL FeOOH with 216 mg of poly(acrylic acid) (PAA, ~1800) in 600 mL of DI water overnight under the magnetic plate. The PAA-modified FeOOH was purified and washed with DI water three times at 11,000 rpm for 15 minutes and dispersed in 12 mL DI water. For the silica coating, 150 μL TEOS was added to a mixture containing 2 mL FeOOH dispersion, 20 mL ethanol, and 250 µL ammonium solution twice with a 1-h interval. The solution was magnetically stirred overnight. Afterward, FeOOH@$SiO_2$ was isolated and washed with ethanol once and water three times at 14,500 rpm for 10 min. Finally, to reduce the FeOOH@$SiO_2$, 25 mL TEG was heated to 280 °C under nitrogen, to which 250 µL concentrated FeOOH@$SiO_2$ was injected. The reaction was kept for 8 h under nitrogen protection. Then final product $Fe_3O_4$@$SiO_2$ was washed with ethanol and water three times and dried under nitrogen for further use.

#### 3. Carboxyl (-COOH) modification of the nanorods

The carboxyl-modified APTES was synthesized by directly mixing 4 g succinic anhydride with 4 mL APTES in 20 mL DMF. After incubating on the magnetic plate overnight, the carboxyl-modified APTES could be used without any purification. Next, 4 mL carboxyl-modified APTES in DMF was added to the $Fe_3O_4$@$SiO_2$ dispersion in DMF (1 mg/mL) and incubated at room temperature on a stirring plate 37 h. After that, the product was isolated by the external magnetic field and washed with DMF and ethanol three times. The obtained magnetic nanorods was dispersed in DI water at a concentration of 10 mg/mL.

#### 4. Antibody modification to form the NanO-stirBar (NOB) for EV isolation

The antibody was conjugated to the magnetic nanorods via EDC/NHS coupling. Briefly, 100 µL 10 mg/mL carboxyl-modified $Fe_3O_4$@$SiO_2$ was added to 1 mL MES (0.1 M, pH 5) solution containing (2 mg/mL EDC, 1 mg/mL NHS). The mixture was incubated at room temperature for 2 h. After that, the activated magnetic nanorods were isolated and washed with 1× PBS three times, and then redissolved in the borate buffer (10 mM sodium borate, 150 mM NaCl, pH 8.5) containing 250 ng/mL primary antibody and incubated at 4 °C overnight. The 0.5% Glycine solution prepared in 1× PBS was used to deactivate the activated carboxyl group on the nanorods after antibody conjugation through overnight incubation at 4 °C. At last, the final EV-targeting NOBs were washed with 1× PBS three times and dispersed in 1× PBS at a concentration of 1 mg/mL.

#### 5. Chip Fabrication

The multi-well chip contains two parts: a PDMS chamber and the cover glass substrate. Briefly, the cover glass (24 × 50 mm, 1.5H) was immersed in piranha solution (98% $H_2SO_4$ and 30% $H_2O_2$ mixed at a volume ratio of 3:1). The solution was heated to boiling for 30 min and cooled down to room temperature. The pretreated cover glass was washed with ethanol five times and dried with nitrogen. The PDMS chamber was obtained by directly pouring the elastomer mixture (Dow Corning Sylgard 184, base and curing reagent w/w = 10:1) into the plastic mold and kept at 55 °C overnight. After curing, the PDMS chamber could be directly peeled and attached to the cover glass by plasma treatment.

#### 6. Circular Probe Preparation

The circular probe was prepared by mixing 5 µL phosphorylated padlock probe (10 µM), 5 µL initiator (10 µM), 4 µL 10× T4 ligase reaction buffer (500 mM Tris-HCl, 100 mM $MgCl_2$, 100 mM DTT, 10 mM ATP), 2 µL T4 ligase ( 40 U/ µL), and 24 µL nuclease-free water, and incubating at 37 °C for 3 h and then at 70 °C for 15 min. After that, 4 µL Exonuclease I, 4 µL Exonuclease III and 2 µL nuclease-free water were added to digest the un-ligated padlock probe and initiator at 37 °C overnight. The reaction was terminated by incubation at 80 °C for 20 min. Finally, the obtained circular probe was stored at 4 °C for further use.

#### 7. RCA reaction

The RCA reaction was performed in a 20 µL solution of 2 µL 10× phi29 buffer (500 mM Tris-HCl, 100 mM $MgCl_2$, 100 mM $(NH_4)_2SO_4$, 40 mM DTT), 0.25 µL phi29 DNA polymerase (10 unit/µL), 1 µL of 0.1 µM circular probe, 1 µL of 0.5 µM initiator, 9.75 µL DI water, 3 µL of 1 mM dNTP, 2 µL 10× SYBR gold. The reaction was added to a 96-well PCR plate and monitored using a Bio-Rad CFX Connect Real-Time PCR Detection System. The reaction mixture was incubated at 37 °C, and fluorescence curves were recorded at 30-s intervals.

#### 8. NOBL-SPA for dual-maker Analysis

The dual-marker detection assay is similar to the single-marker assay. Briefly, after fixing the captured EV, a mixture of 1 µL 0.05 µM circular probe, 1 µL 0.25 µM recognition probe, and 8 µL 1× PBS was added to recognize the target molecule (protein and miRNA) at room temperature for 30 min on the magnetic stirring plate (360 rpm). After removing the supernatant and washing the captured EV, the RCA reaction buffer (1 µL 10× phi29, 1 µL of 0.125 mg/mL BSA, 1 µL of 1 mM dNTP, 1 µL phi29 DNA polymerase (2.5 Units/mL), and 9 µL DI water ) was added to the well and incubate at 37 °C for 30 min, which was followed by addition of 1 µL of 2.5 µM detection probe (Alexa 488 for miRNA and Alexa 647 for surface protein) and incubation for 30 min. After being washed with 1× PBS three times and dispersed in 10 µL 1× PBS, the sample was ready to be imaged.

#### 9. Detection in Cell Culture Media and Clinical Samples

The cell culture medium was utilized directly without any pretreatment. Five µL cell culture medium and 5 µL 1× PBS were added to the well containing the NOB for NOBEL-SPA. The serum samples were firstly pre-treated with ultracentrifugation at 10,000 *g* for 10 min to remove the cell debris; and then 1 µL serum and 9 µL 1× PBS (for Exo-NOBs), or 5 µL serum and 5 µL 1× PBS (for the NOBs targeting other markers) were added to the well for NOBEL-SPA.

#### 10. Enzyme-linked immunosorbent assay (ELISA)

The 96-well plate was firstly modified with anti-CD9, anti-CD63, and anti-CD81 antibodies by incubating with 100 µL of 2 µg/mL anti-CD9, anti-CD63, and anti-CD81 antibodies at 4 °C overnight. After washing with 1× PBS twice, 300 µL 1% BSA in 1× PBS was employed to block the plate at room temperature for 2 hours. Then 100 µL EV solution in 1× PBS containing 1% BSA was added and incubated at room temperature for 2 h and subsequently at 4 °C for 6 hours. After supernatant removal and two washes with 1× PBS containing 1% BSA, 100 µL of 1 µg/mL biotinylated anti-CD63 was added and incubated at 4 °C overnight. After another round of wash, the plate was incubated with 100 µL of 0.05 µg/mL streptavidin-modified horseradish peroxidase (HRP) at room temperature for 30 min and subsequently incubated with the SuperSignal™ West Pico PLUS Chemiluminescent Substrate for signal development. The signal was read at BioTek Synergy H1 Hybrid Multi-Mode Microplate Reader.

#### 11. Western Blot

For WB, the EVs were lysed directly with the SDS loading buffer containing 10% SDS, 500 mM DTT, 50% Glycerol, 500 mM Tris-HCL, and 0.05% bromophenol blue dye and separated on a 15% SDS-PAGE gel. The gel was transferred onto a nitrocellulose membrane (BIO-RAD). Membranes were blocked for 1 h at room temperature with 5% BSA (RPI, Research Products International) in the TBST (1×Tris-Buffered Saline containing 0.1% Tween 20 detergent) solution and followed by incubation with the primary antibody solution overnight at 4 °C. The following antibodies were used at 1:500 dilution: anti-CD63 (Novus Biologicals) and anti-CD81 (Sigma-Aldrich). Then, the membrane was incubated with the HRP-conjugated goat-anti-mouse IgG (Invitrogen, 1:20,000 dilution) for 1 h and subsequently with the chemiluminescent substrate before being imaged by Odyssey XF Imaging System (LI-COR Biosciences).

##### 12. EV miRNA quantification via RT-qPCR

The enclosed miRNA in EV was firstly isolated by the miRNeasy Mini Kit (QIAGEN) and quantified by RT-qPCR using the TaqMan Probe primers and the TaqMan MicroRNA Reverse Transcription Kit (Thermofisher Scientific). The reverse transcription mix was composed of 1.1 µL RNase-free water, 1.0 µL 10× reverse transcriptase buffer, 0.13 µL RNase inhibitor (20 U/µL), 0.1 µL dNTP mix (100 mM), 0.67 µL Multiscribe RT enzyme (50 U/µL), and 2.0 µL 5× RT primer. 5 µL of the RT mix and 5 µL of the sample were added to individual tubes and underwent RT in a Bio-Rad CFX thermocycler. The reaction protocol was as follows: 16 °C for 30 min, 42 °C for 32 min, and 85 °C for 5 min (3 cycles). Following RT, qPCR was carried out. The reaction mixture contained 1 µL of the RT product, 0.1 µL DMSO, 1.0 µL ethylene glycol (> 99%), 0.5 µL magnesium chloride (25 mM), 4.9 µL RNase-free water, 2 µL Taq 5x master mix (NEB), and 0.5 µL TaqMan probes. The reaction protocol was as follows: 95 °C for 3min, followed by a denaturing step at 95 °C for 20 s and a combined annealing and extension step at 60 °C for 30 s and at 68 °C for 1 min that cycled 40 times.

##### 13. Nanoparticle Tracking Analysis (NTA) Characterization

NTA was carried out by NanoSight NS300 (Malvern Instruments) equipped with a low-volume flow cell manifold and a 405 nm laser module. Three videos with 30 same samples were taken at a rate of 25 frame/s.

##### 14. Scanning Electron Microscope (SEM) Sample Preparation and Characterization

To image the NOB, 10 µL of 10 ng/mL NSB solution was directly dropped on the silica wafer and dried in the air.

For EV imaging, the silica wafer was firstly cleaned with piranha solution at 88 °C for 10 min. After being washed with ethanol three times, the pretreated silica wafer was sequentially functionalized with primary amine and aldehyde by dropping 5% APTES in ethanol (15 min) and 1% glutaraldehyde in 1× PBS on the surface (1 h). After that, 10 µL of the EV solution was dropped on the aldehyde-modified silica surface and incubated at room temperature for 1 h. And then, the residual aldehyde groups were deactivated with the TEA buffer (0.1 M Tris, 50 mM ethanolamine pH 9) and 0.05% w/v casein in 1× PBS. Each treatment lasted for 30 min. After being washed with 1× PBS three times, the immobilized EVs were fixed with 0.1% glutaraldehyde and 2% paraformaldehyde in 1× PBS for 30 min and then dehydrated with 20%, 40%, 60%, 80%, and 100% ethanol sequentially. Finally, the sample was dried in the air at room temperature.

To image the EV captured on the NOBs, the sample was fixed with 0.1% glutaraldehyde and 2% paraformaldehyde in 1× PBS for 30 min; then dehydrated with 20%, 40%, 60%, 80%, and 100% ethanol sequentially; and at last dispersed in DI water before being dropped on the silica wafer and dried in the air.

Before SEM imaging, all of the sample-containing silica wafers were coated with a cPt/Pd sputter coater. The SEM was performed with an NNS450 instrument (ThermoFisher Scientific).

#### Supplemental Table

**Table S1. Recent single EV detection method comparison.**

| Leading Author & Year of Publication (Cited as Ref. 16 in Main Text) | Detection Technique | Limit of Detection; Sample Volume | Assay Time | Single EV Capture | Directly work with biofluids? |
|---|---|---|---|---|---|
| **Surface Protein** | | | | | |
| H. Im et al, 2023 | CFM | 70 particles/μL; 30 μL | 2.5 h | Specific surface structure | No |
| T. Y. Hu et al, 2022 | Dark field microscopy/scatter light | 39 particle/μL; 1 μL | > 4 h | Dilute EV sample | Yes |
| D. A. Issadore et al, 2020 | Simoa® | 9 particles/μL; 10 μL | Not reported | Control bead : EV ratio | Yes |
| **miRNA or mRNA** | | | | | |
| H.-W. Li et al, 2019 | TIRFM | 378 copies/μL; 10 μL | 2 h | Dilute EV sample | Yes |
| **mRNA/miRNA and Protein** | | | | | |
| C. Bai et al, 2020 | TIRFM | Not mentioned; 90 μL | 6 h | Dilute EV sample | No |
| NOBEL-SEA (reported in the present work) | CFM | 3-4 particles/μL; < 5 μL | ~ 4 h | Unique features of NOB | Yes |

#### Supplementary Figures

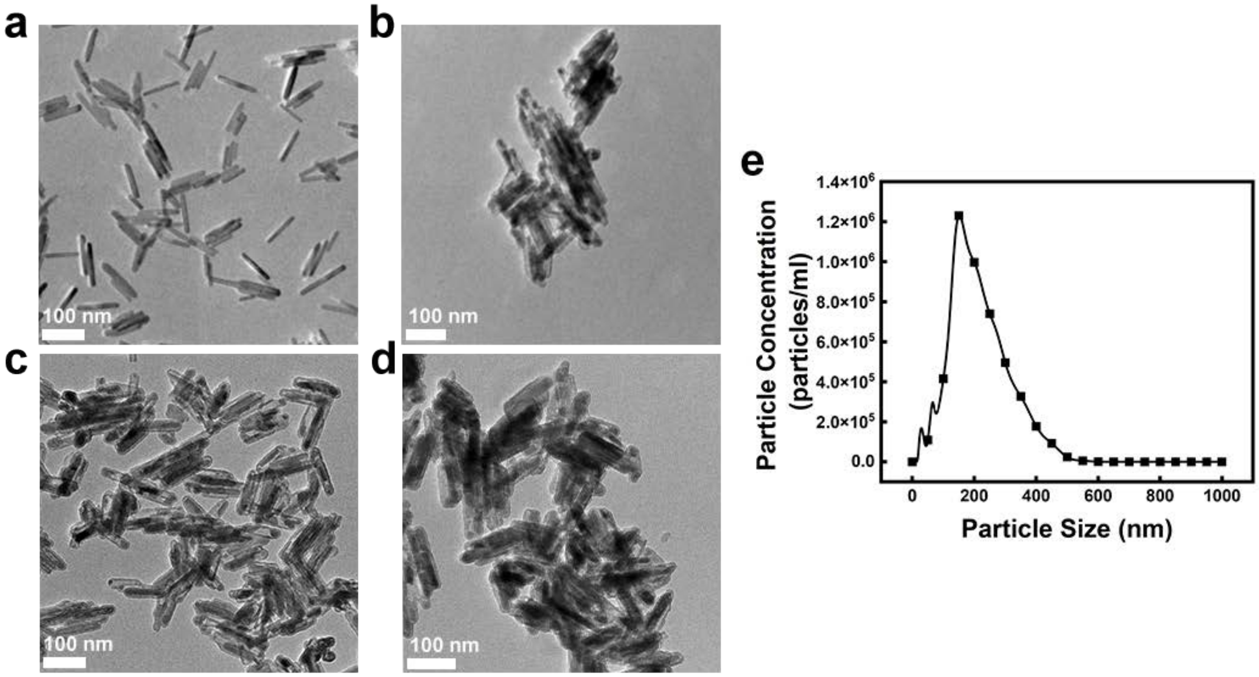


**Figure S1. TEM images of the NSB at different stages of synthesis and modification**. The hydroxy iron oxide(FeOOH) nanorod was firstly produced (a), then reduced into the magnetic $Fe_3O_4$ by trimethylene glycol (TEG), coated by silica (b), then modified with –COOH groups through APTES and succinic anhydride (c). At last, the nanorods were conjugated with antibodies to form the EV-targeting NOBs (d), the size distribution of which was collected by NTA (e).

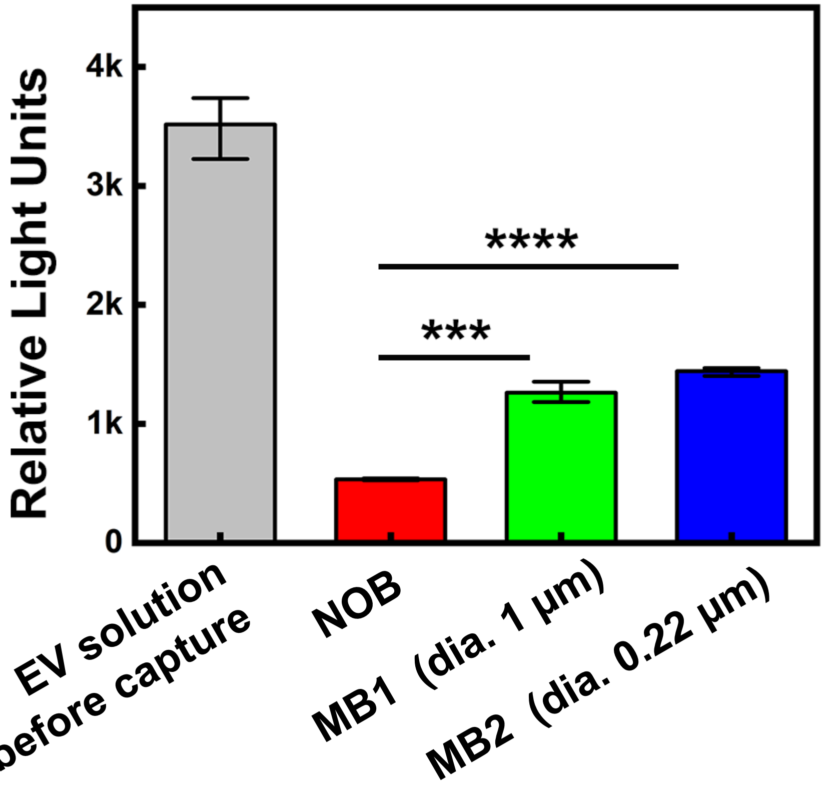


**Figure S2. Exosome capture efficiency comparison between the NSBs and the commercial magnetic beads with diameters (dia.) around 1 and 0.22 μm**. The evaluation was performed by detecting the presence of the exosomal marker CD63 in the supernatant with ELISA after exosome capture by the antibody-conjugated beads. The Relative Light Units shown in the Y-axis was the signal from ELISA.

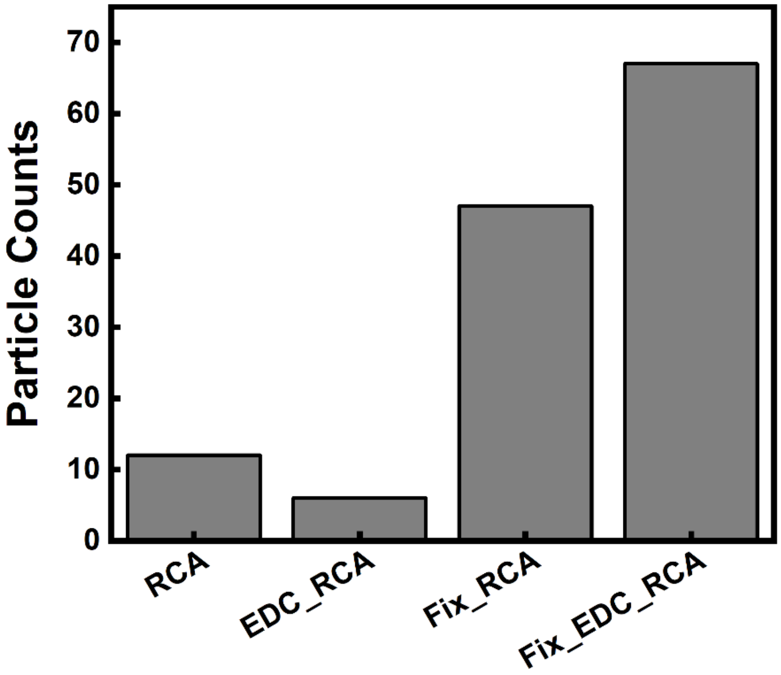


**Figure S3. The necessity of EDC crosslink and PFA fixation for detection of EV miRNAs**. The fixation and crosslinking conditions greatly increased the number of fluorescent particles counted in CFM through DNF labeling upon recognizing the target miRNA miR-155.

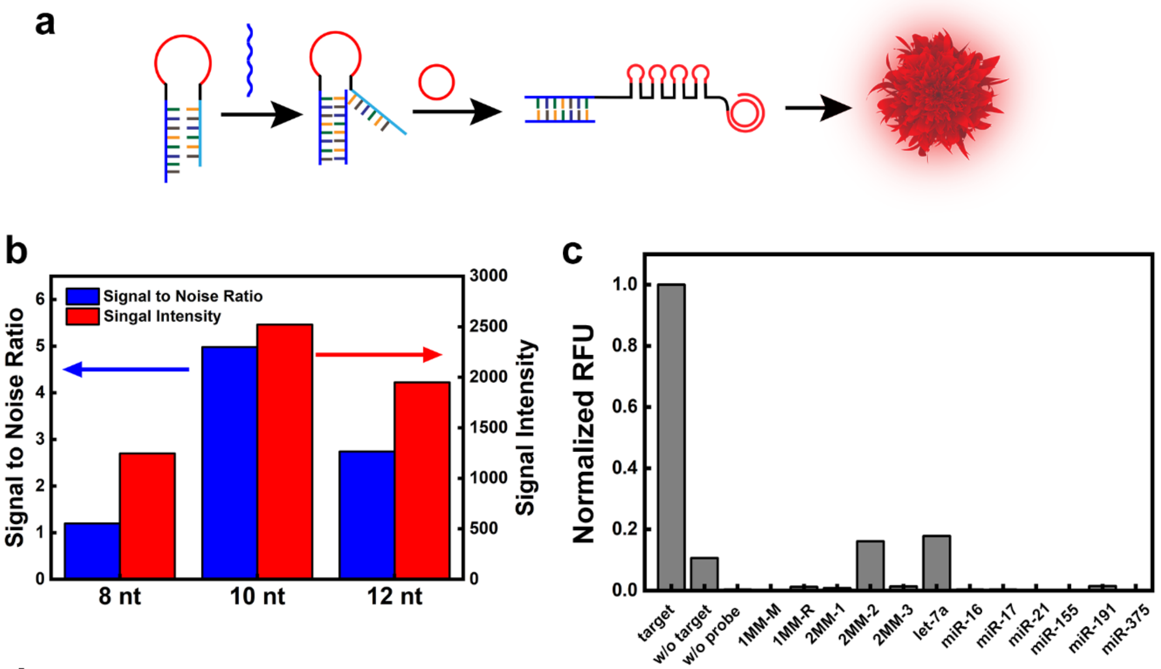


d

| Name | Sequence (5'-3') |
|---|---|
| Padlock probe | /5Phos/G ACT AAC GAC AAC G CGTG TTA GCA A GCG ATA CGC G TA TCG CTA TGG CAT ATC GTA CGA TAT GCC TGC TAA CAC GC C GAA TCC TA |
| Initiator | CGT TGT CGT TAG TCT AGG ATT CGG CGT GTT AGC A |
| Let-7a | UGA GGU AGU AGG UUG UAU AGU U |
| miR-16 | UAGCAGCACGUAAAUAUUGGCG |
| miR-17 | CAAAGUGCUUACAGUGCAGGUAG |
| miR-21 | UAG CUU AUC AGA CUG AUG UUG A |
| miR-122 | UGG AGU GUG ACA AUG GUG UUU G |
| miR-155 | UU AAU GCU AAU CGU GAU AGG GGU |
| miR-155 Recognition Probe (8 nt stem) | ACC CCT ATC ACG ATT AGC ATT AA G TCT CCC A GT CTA GGA TTC GGC GTG TTA GCA TTA ATG CT |
| miR-155 Recognition Probe (10 nt stem) | ACC CCT ATC ACG ATT AGC ATT AA G TCT CCC A GT CTA GGA TTC GGC GTG TTA GCA TTA ATG CTA A |
| miR-155 Recognition Probe (12 nt stem) | ACC CCT ATC ACG ATT AGC ATT AA G TCT CCC A GT CTA GGA TTC GGC GTG TTA GCA TTA ATG CTA ATC |
| miR-155 one base mismatch-M (1MM-M) | TT AAT GCT AAT CCT GAT AGG GGT |
| miR-155 one base mismatch-R (1MM-R) | TT AAT GCT AAT CGT GAT ACG GGT |
| miR-155 two bases mismatch-1 (2MM-1) | TT AAT ACT AAT CCT GAT AGG GGT |
| miR-155 two bases mismatch-2 (2MM-2) | TT AAT ACT AAT CGT GAT ACG GGT |
| miR-155 two bases mismatch-3 (2MM-3) | TT AAT GCT AAT CCT GAT ACG GGT |

**Figure S4**. a) Scheme of miRNA recognition by a hairpin probe. b) Optimization of the size of the stem region of the hairpin probe. The fluorescent signal was obtained by monitoring the RCA in a real-time PCR instrument. The signal intensity was the maximum reading (in RFU) after 2 hours of reaction, with the signal-to-noise ratio being the ratio between the positive reading and the negative reading. c) Specificity of miR-155 detection. All signals were obtained as done in b). The Y-axis represents the normalized RFU obtained by dividing the signal intensity in the presence of the target or any of the mutated or non-target strand by that acquired without (w/o) the target. d) Summary table for the sequences used for miR-155 detection, stem length optimization, and specificity test. The sequence in blue is complementary to miR-155; that in red is the initiator that could recognize the circular probe and trigger RCA; and the underlined sequence is the stem region of the hairpin probe.

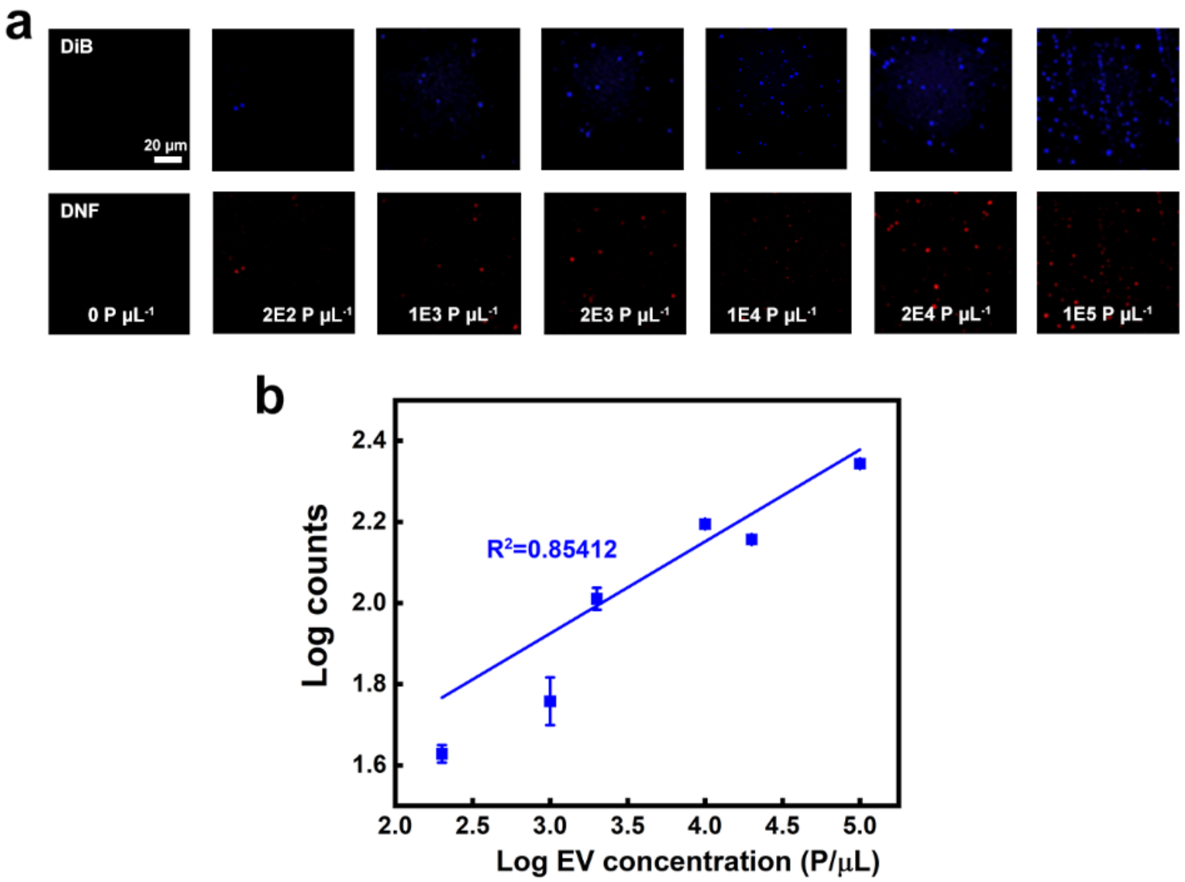


**Figure S5. miR-155 detection performance.** a) Representative confocal images of the stained EV (blue) and the DNFs for labeling miR-155 obtained with different standard EV concentrations; b) The calibration curve plotting the logarithmic value (Log) of the EV counts vs. the Log value of the input EV concentration.

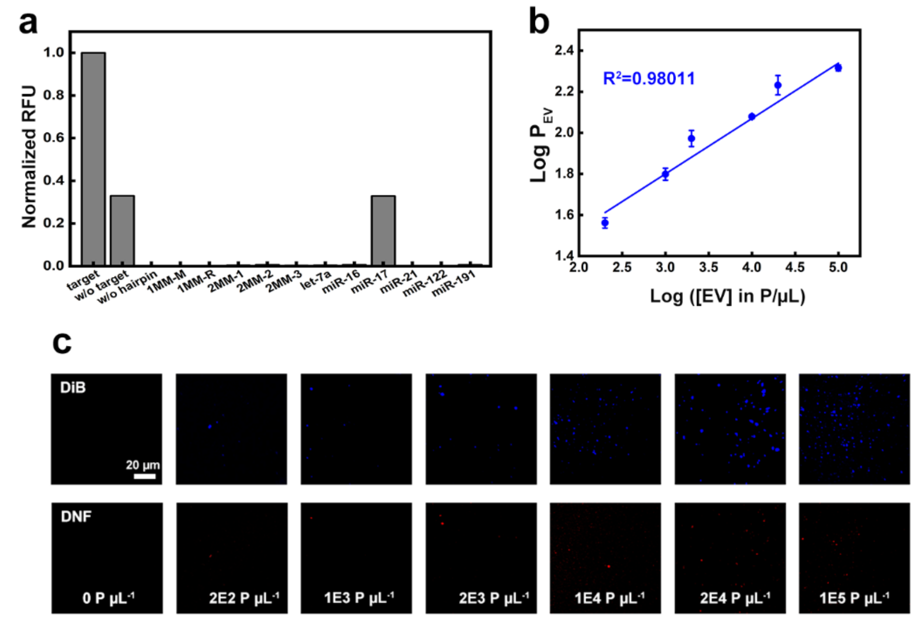


d

| Name | Sequence (5'-3') |
|---|---|
| Padlock probe | /5Phos/G ACT AAC GAC AAC G CGTG TTA GCA A GCG ATA CGC G TA TCG CTA TGG CAT ATC GTA CGA TAT GCC TGC TAA CAC GC C GAA TCC TA |
| miR-21 Recognition Probe | TCA ACA TCA GTC TGA TAA GCT A GT GTG TGA GTC TAG GAT TCG GCG TGT TAG CA T AGC TTA TCA |
| miR-122 one base mismatch-L (1MM-L) | TGG ACT GTG ACA ATG GTG TTT G |
| miR-122 one base mismatch-R (1MM-R) | TGG AGT GTG ACA ATG GTC TTT G |
| miR-122 two bases mismatch-1 (2MM-1) | TGG ACT GTG ATA ATG GTG TTT G |
| miR-122 two bases mismatch-2 (2MM-2) | TGG ACT GTG ACA ATG GTC TTT G |
| Let-7a | UGA GGU AGU AGG UUG UAU AGU U |
| miR-16 | UAGCAGCACGUAAAUAUUGGCG |
| miR-17 | CAAAGUGCUUACAGUGCAGGUAG |
| miR-21 | UAG CUU AUC AGA CUG AUG UUG A |
| miR-122 | UGG AGU GUG ACA AUG GUG UUU G |
| miR-155 | UU AAU GCU AAU CGU GAU AGG GGU |

**Figure S6. miR-122 detection performance via NOBEL-SPA.** a) miR-122 detection specificity test. The signal was obtained from the real-time PCR instrument by monitoring the fluorescent signal change which was normalized against the signal intensity obtained with miR-122. b) The calibration curve plotting the Log value of the EV count vs. the Log value of the input EV concentration. c) Confocal images of the stained EVs (blue) and the DNFs labeling miR-122 obtained with different standard EV concentrations. d) The table summarizing the sequences employed in miR-122 detection and specificity test. The sequence in blue is complementary to miR-122; that in red is the initiator that could recognize the circular probe and trigger RCA; and the underlined sequence is the stem region of the hairpin probe.

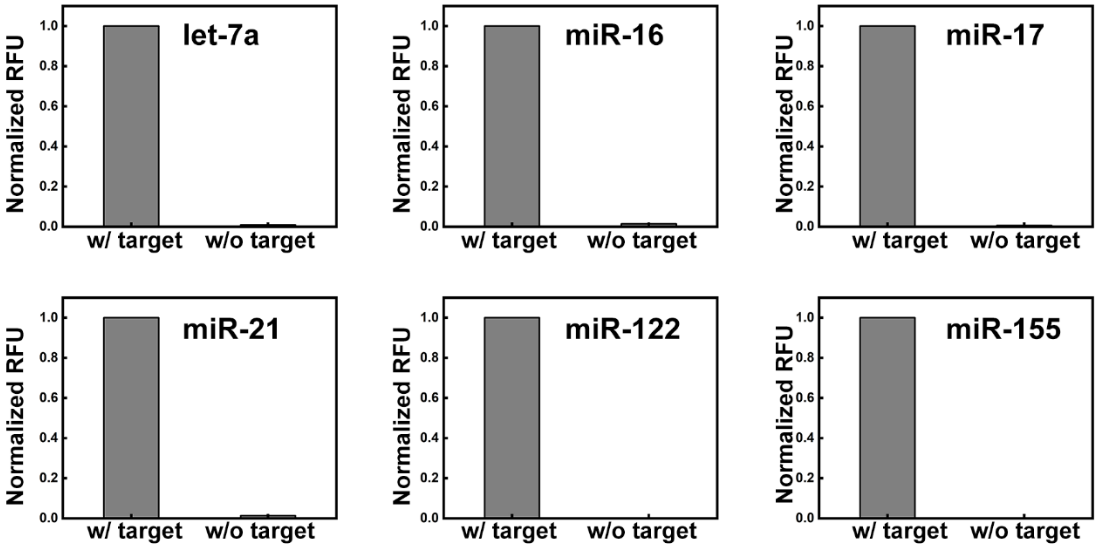


| Name | Sequence (5'-3') |
|---|---|
| Padlock probe | /5Phos/G ACT AAC GAC AAC G CGTG TTA GCA A GCG ATA CGC G TA TCG CTA TGG CAT ATC GTA CGA TAT GCC TGC TAA CAC GC C GAA TCC TA |
| Let-7a | UGA GGU AGU AGG UUG UAU AGU U |
| Let-7a Recognition Probe | AAC TAT ACA ACC TAC TAC CTC A GT GTG TGA GTC TAG GAT TCG GCG TGT TAG CA T GAG GTA GTA |
| miR-16 | UAGCAGCACGUAAAUAUUGGCG |
| miR-16 Recognition Probe | C GCC AAT ATT TAC GTG CTG CTA GT GTG TGA GTC TAG GAT TCG GCG TGT TAG CA TAG CAG CAC G |
| miR-17 | CAAAGUGCUUACAGUGCAGGUAG |
| miR-17 Recognition Probe | CT ACC TGC ACT GT A AGC ACT TTG GT GTG TGA GTC TAG GAT TCG GCG TGT TAG CA CAA AGT GCT T |
| miR-21 | UAG CUU AUC AGA CUG AUG UUG A |
| miR-21 Recognition Probe | TCA ACA TCA GTC TGA TAA GCT A GT GTG TGA GTC TAG GAT TCG GCG TGT TAG CA T AGC TTA TCA |
| miR-122 | UGG AGU GUG ACA AUG GUG UUU G |
| miR-122 Recognition Probe | CAA ACA CCA TTG TCA CAC TCC A GT GTG TGA GTC TAG GAT TCG GCG TGT TAG CA T GGA GTG TGA |
| miR-155 | UU AAU GCU AAU CGU GAU AGG GGU |
| miR-155 Recognition Probe | ACC CCT ATC ACG ATT AGC ATT AA G TCT CCC A GT CTA GGA TTC GGC GTG TTA GCA TTA ATG CTA A |

**Figure S7. Detection of multiple miRNA targets.** a) Feasibility test for detection of multiple miRNA targets using the hairpin probes listed in the summary table together with the target sequences and the padlock probe sequence. The concentration of the target strand, the recognition hairpin probe, and the circular probe was 0.05 µM, 0.025 µM, and 0.005 µM, respectively. The reaction was carried out and monitored by the Bio-Rad CFX Connect Real-Time PCR Detection System. After 2 hours of reaction, the RFU was normalized against that with the target for each strand. In the sequence summary table, the sequence in blue is complementary to the target miRNA; that in red is the initiator that could recognize the circular probe and trigger RCA; and the underlined sequence is the stem region of the hairpin probe.

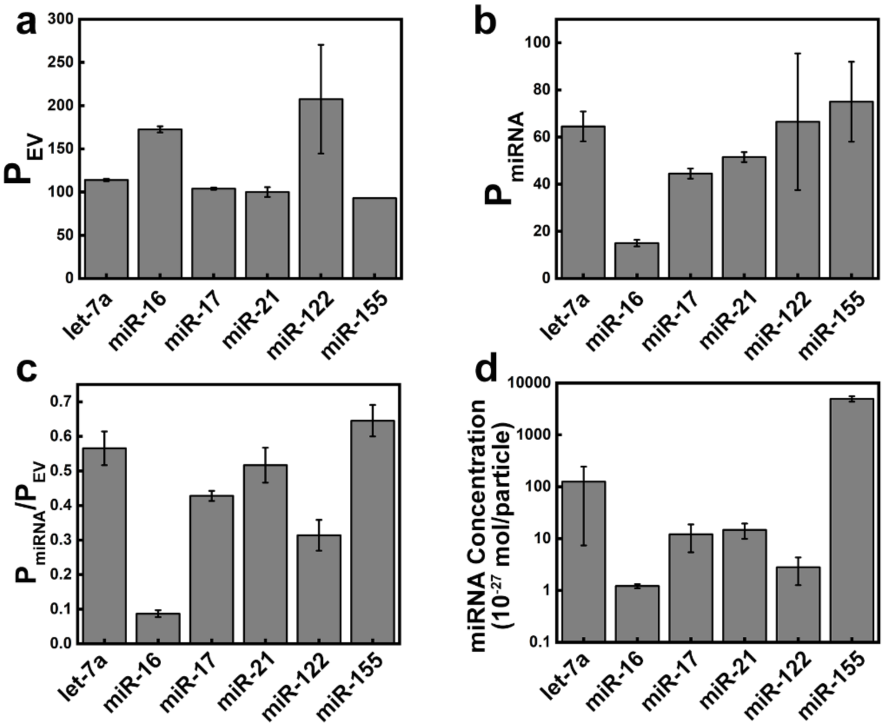


**Figure S8. miRNA profile in the standard EVs derived from COLO1 cells as indicated by the manufacturer.** a-c) The bar plots of the counts of the stained EVs ($P_{EV}$) (a) and the EVs labeled by DNFs via recognition of different miRNA targets (b), as well as the ratios of $P_{miRNA}/P_{EV}$, obtained with NOBEL-SPA using an input EV of $10^5$ EV particles. d) The miRNA concentrations expressed in mole of miRNA per EV particle using the miRNA concentration obtained by RT-qPCR and the EV counts given by NTA. The total RNA in $10^8$ EVs were extracted by the miRNeasy Mini Kit, which yielded a solution of 40 µL. Each RT-qPCR reaction took 5 µL of the extracted miRNA.

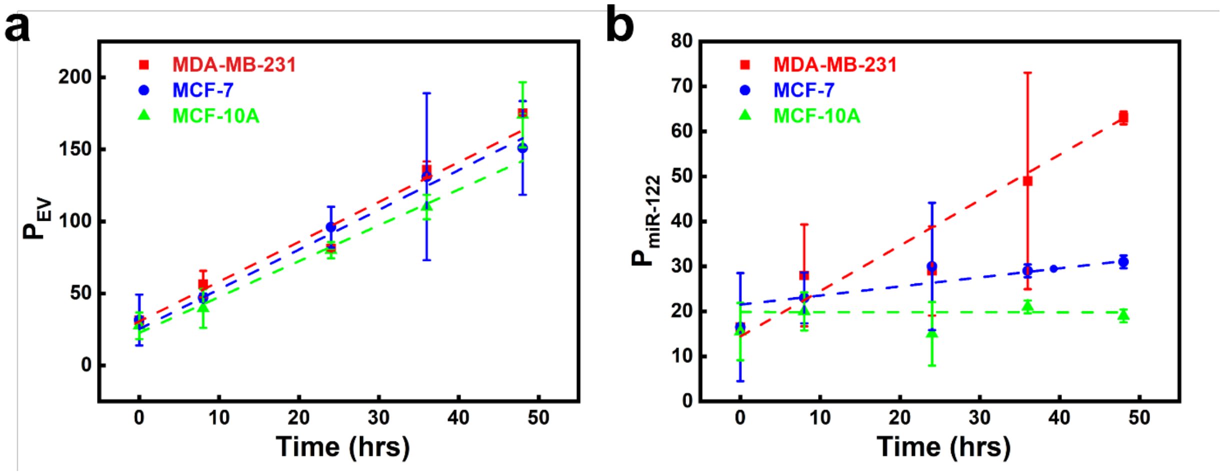


**Figure S9. Monitoring EV secretion in three breast cell line (MCF-10A, MCF-7, and MDA-MB-231) cell culture media by NOBEL-SPA.** a) $P_{EV}$ and b) $P_{miR-122}$ obtained by NOBEL-SPA in the culture media of MCF-10A, MCF-7, and MDA-MB-231 cells collected at different time points after switching to the EV-free medium. Two repeated measurements were carried out for each sample, i.e. 10 µL of the culture medium from one cell line collected at one specific time point. In each repeat, a total of 10 images were taken. The total particle counts found in these 10 images were averaged for the two repeats, and reported in the plots.

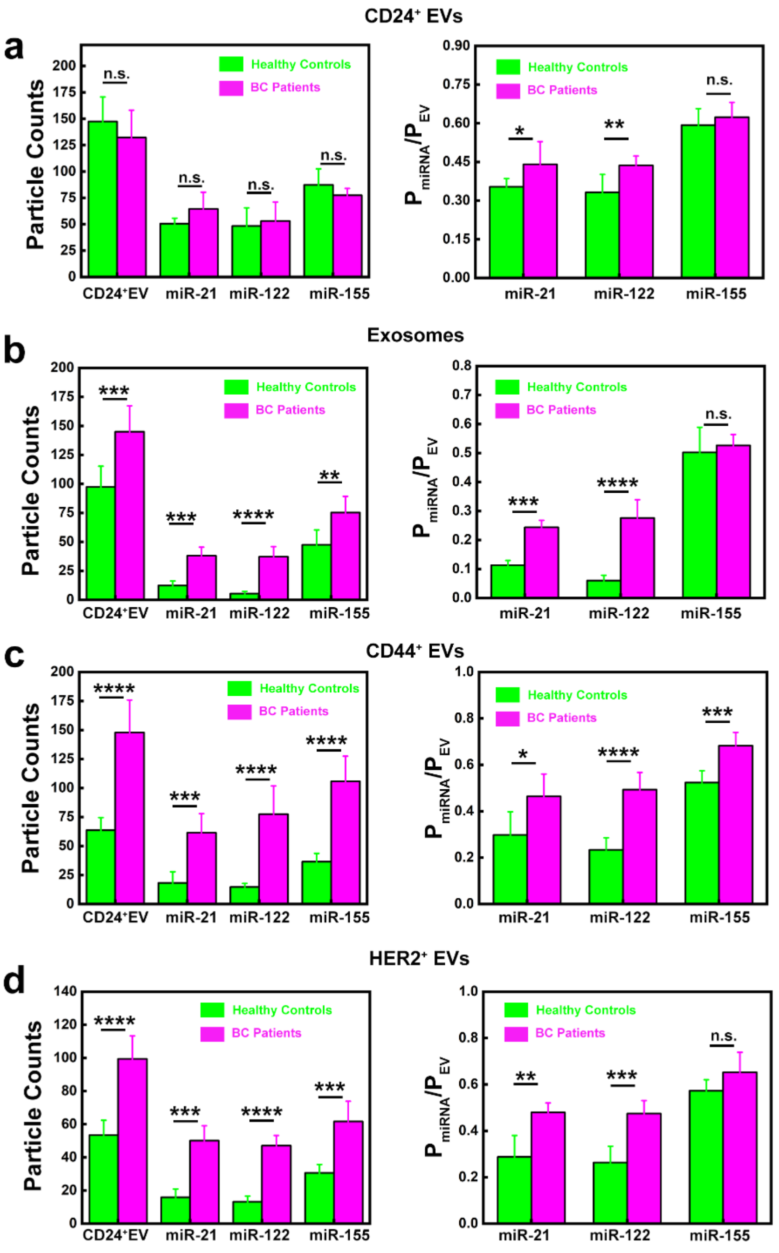


**Figure S10. Detection of exosomes or the EVs having the surface protein of CD44, CD24, or HER2, and carrying specific miRNAs in clinical samples by the single-marker NOBEL-SPA.** The bar plots in a-d) compared the particle counts ($P_{EV}$ or $P_{miRNA}$) (in the left panel) and the ratios of $P_{miRNA}/P_{EV}$ (in the right panel) obtained from the CD24$^{+}$ EVs (a), the exosomes (b), the CD44$^{+}$ EVs (c), and the HER2$^{+}$ EVs (d) between healthy controls (green) and BC patients (magenta). Two repeated measurements were carried out for each sample, i.e. 1 μL (for exosome) or 5 μL (for other EVs) of the human serum from one individual. In each measurement, a total of 10 images were taken. The sum of the particle counts found in the 10 images, or the average ratio of the particle counts of the 10 images collected in each repeated measurement, was used to calculate the average and standard deviation (error bar) of the two repeats from each individual. Then the average and standard deviation of each cohort (n = 3) were reported in the plots.

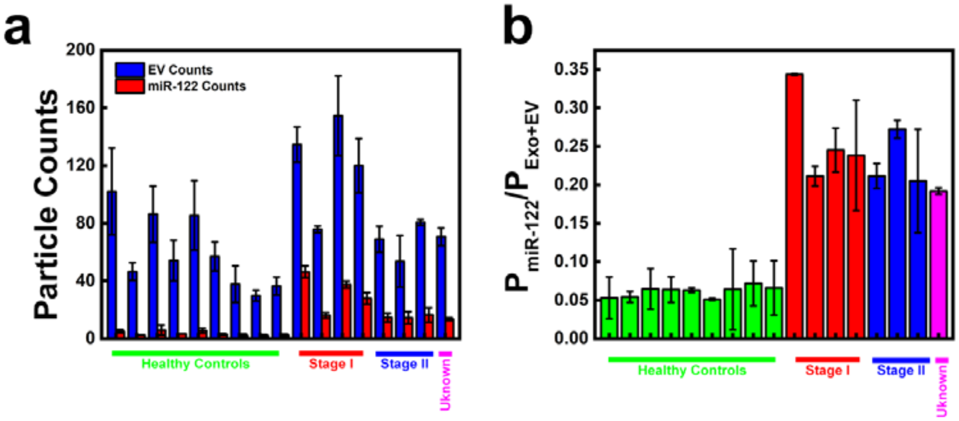


**Figure S11. Detection of exosomes carrying miR-122 in clinical samples by the single-marker NOBEL-SPA.** Two repeated measurements were carried out for each sample, i.e. 1 μL of the human serum from one individual. In each measurement, a total of 10 images were taken. The sum of the particle counts found in the 10 images, or the average ratio of the particle counts of the 10 images collected in each repeated measurement, was used to calculate the average and standard deviation (error bar) of the two repeats from each individual reported in the plots.

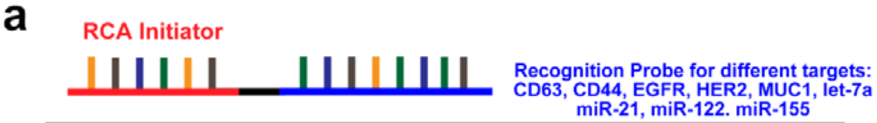


| Name | Sequence (5'-3') |
|---|---|
| Protein Detection Padlock Probe (R3CP) | /5Phos/CAC AGA TAC GTT ATA CCC GGT CGC TTT AGC CTT CTT ACT GAG |
| Protein Detection Initiator | CG TAT CTG TGC TCA GTA AGA |
| CD63 Recognition Probe (R3CD63RP) | CAC CCC ACC TCG CTC CCG TGA CAC TAA TGC TA T TTT T CG TAT CTG TGC TCA GTA AGA |
| CD44 Recognition Probe (R3CD44RP) | CCA AGG CCT GCA AGG GAA CCA AGG ACA CAG T TTT TCG TAT CTG TGC TCA GTA AGA |
| HER2 Recognition Probe (R3HER2RP) | GGG CCG TCG AAC ACG AGC ATG GTG CGT GGA CCT AGG ATG ACC TGA GTA CTG TCC T TTT T CG TAT CTG TGC TCA GTA AGA |
| EGFR Recognition Probe (R3EGFRRP) | TAC CAG TGC GAT GCT CAG TGC CGT TTC TTC TCT TTC GCT TTT TTT GCT TTT GAG CAT GCT GAC GCA TTC GGT TGA C T TTT CG TAT CTG TGC TCA GTA AGA |
| MUC1 Recognition Probe (R3MUC1RP) | GCA GTT GAT CCT TTG GAT ACC CTG G T TTT TCG TAT CTG TGC TCA GTA AGA |
| Protein Detection Probe | /5Alex647N/TT ATA CCC GGT CGC TTT AGC CT |
| miRNA Detection Padlock Probe (R4CP) | /5Phos/TAG TAT GCG ATG TCA TTA CGA TCA TAA GTT AGG TCC AGA AGC |
| miRNA Detection Initiator | CGC ATA CTA GCT TCT GGA C |
| Let-7a Recognition Probe (R47aRP) | AAC TAT ACA ACC TAC TAC CTC A TTT TTT CGC ATA CTA GCT TCT GGA C |
| miR-21 Recognition Probe (R421RP) | TCA ACA TCA GTC TGA TAA GCT A TTT TTT CGC ATA CTA GCT TCT GGA C |
| miR-122 Recognition Probe (R4122RP) | CAA ACA CCA TTG TCA CAC TCC A TTT TTT CGC ATA CTA GCT TCT GGA C |
| miR-155 Recognition Probe (R4155RP) | ACC CCT ATC ACG ATT AGC ATT AA TTT TTT CGC ATA CTA GCT TCT GGA C |
| miRNA Detection Probe | /5Alex488N/ TGT CAT TAC GAT CAT AAG TTA G |

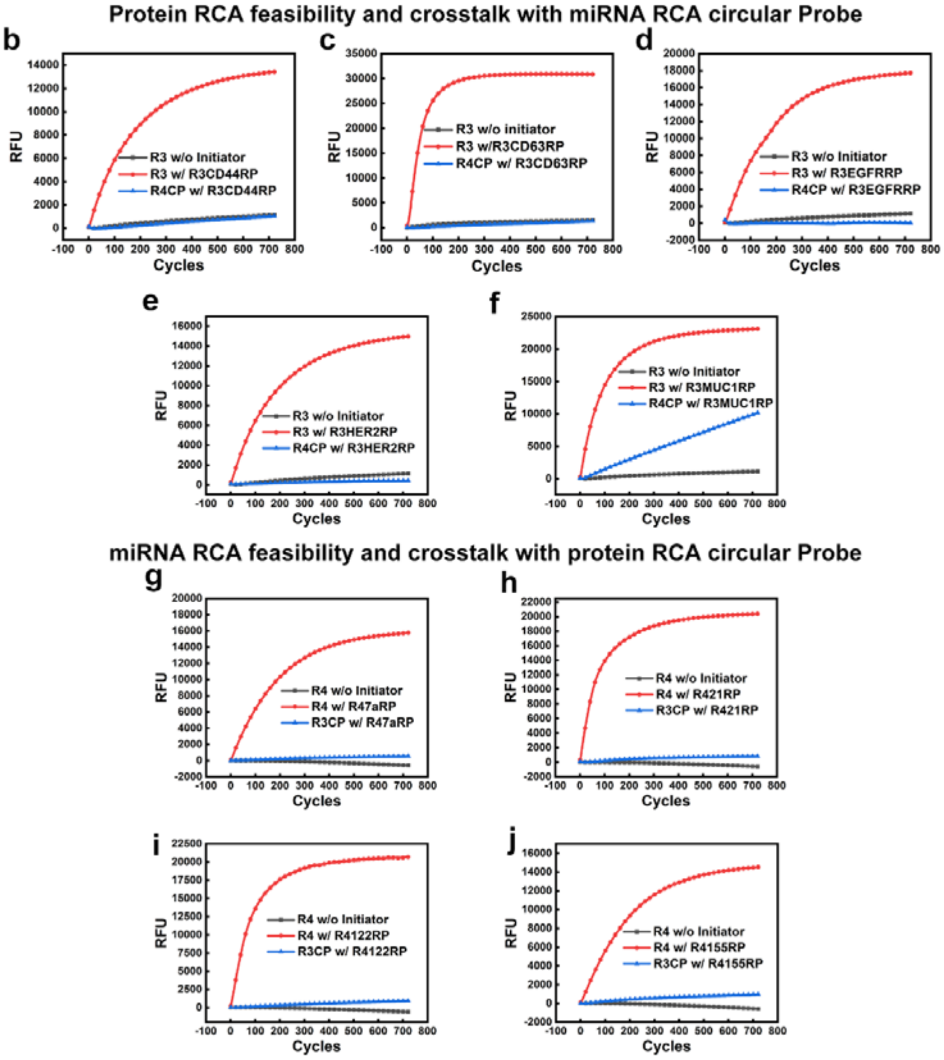


**Figure S12. RCA design and validation for dual-marker detection.** a) The scheme for the design of the target recognition probe and dual-marker detection by NOBEL-SPA. The sequence in blue is the initiator that can recognize the circular probe and initiate RCA; and the sequence in red is for target recognition, which is either the aptamer recognizing the protein target or the complementary strand to the target miRNA. b-f) protein RCA feasibility and crosstalk examination with the miRNA RCA circular probe: b) the probe for CD44 (b), CD63 (c), EGFR (d), HER2 (e), MUC1 (f). g-j) miRNA RCA feasibility and crosstalk examination with the protein RCA circular probe: let-7a (g), miR-21 (h), miR-122 (i), and miR-155 (j). The concentration of the target, the recognition probe and the circular probe was 0.05 µM, 0.025 µM, and 0.005 µM, respectively. The reaction was carried out and monitored by the Bio-Rad CFX Connect Real-Time PCR Detection System.

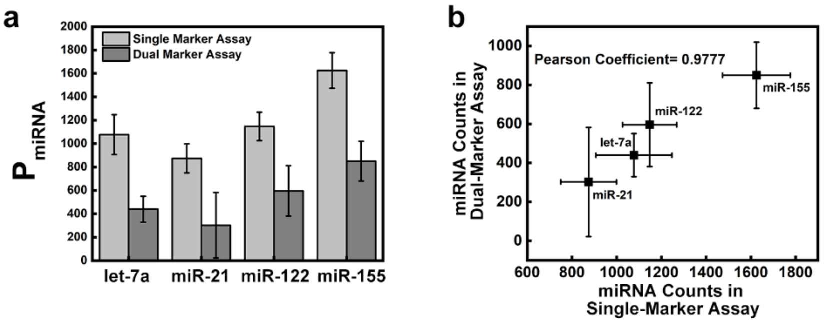


c

| Markers Combination | Pearson Correlation Coefficient |
|---|---|
| CD63/let-7a | 0.1323 |
| CD63/miR-21 | 0.4122 |
| CD63/miR-122 | 0.3677 |
| CD63/miR-155 | 0.4811 |

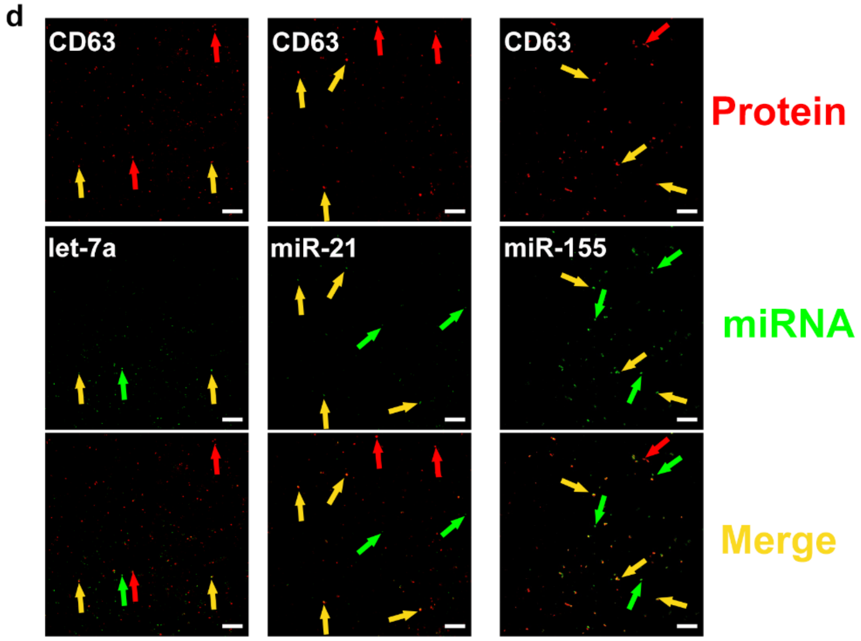


**Figure S13. Feasibility test of dual-marker NOBEL-SPA using the standard EVs.** a) The particle counts from labeling the specific miRNA on the EVs ($P_{miRNA}$) obtained by the single- and dual-marker assay. b) The particle count correlation between the single- and the dual-marker assays. c) The colocalization correlation coefficients for the two fluorescence signals labeling the CD63 and one of the miRNA markers. Two repeated measurements were done for each sample. In each measurement, $10^6$ EV particles were used, 10 images were taken, and the total particle counts in 10 images were obtained. The value and error bar in a-b) represent the average and standard deviations of the total particle counts of the two repeats. d) Representative CFM images of dual-marker NOBEL-SPA, with the red particles representing the protein labels, and the green ones with the miRNA labels. The red, green, and yellow arrows point towards the particles emitting signals from only CD63, only miRNA, or both.

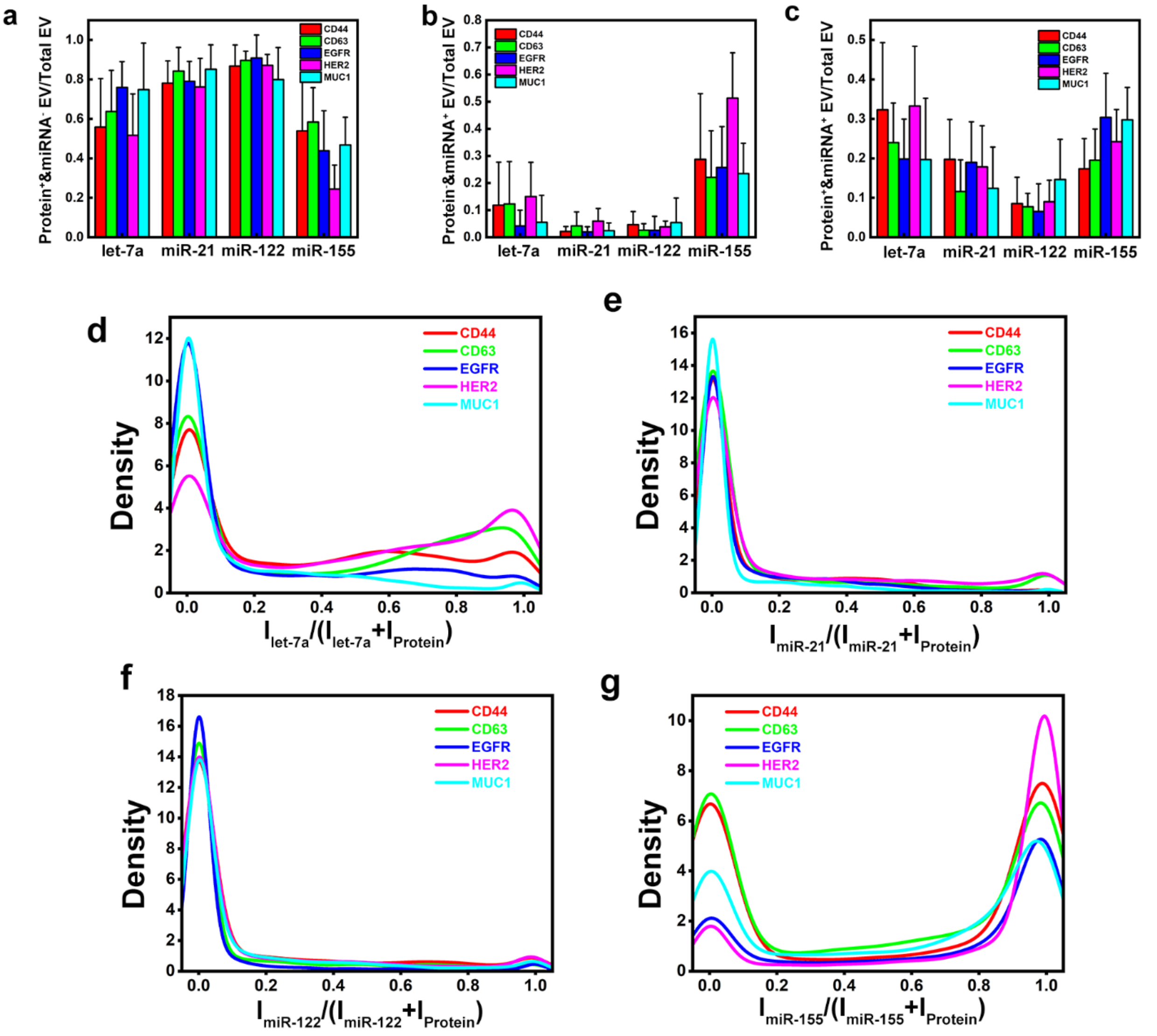


**Figure S14. Results of dual-marker NOBEL-SPA on the A549 -derived EVs**. a-c) The bar plots of the fractions of the EV counts for the protein-only (Protein$^{+}$& miRNA$^{-}$), the miRNA-only (Protein$^{-}$& miRNA$^{+}$), and the dual-marker (Protein$^{+}$&miRNA$^{+}$) EVs among the total EVs detected. A total of 20 protein/miRNA combinations were examined, and 10 images were taken, with the average counts and standard deviations reported in the plots. d-g) The density distribution profiles of the fluorescence intensity ratios of all individual particles detected in the images for the four miRNA markers: d) -- let-7a; e) -- miR-21; f) -- miR-122; and g) -- miR-155. The ratio was calculated by dividing the miRNA signal with the sum intensity of the both the protein and miRNA signals.

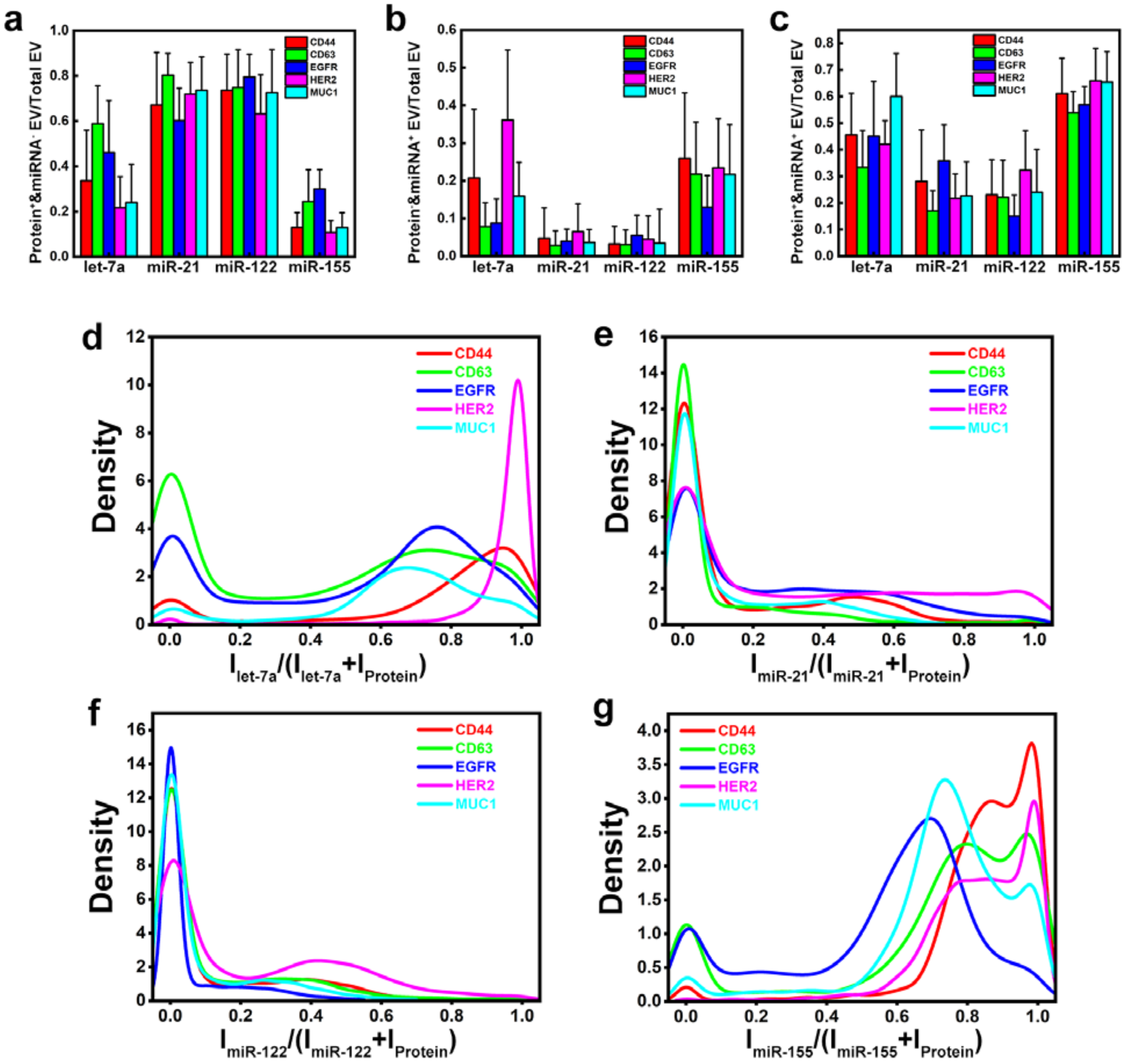


**Figure S15. Results of dual-marker NOBEL-SPA on the HeLa -derived EVs**. a-c) The bar plots of the fractions of the EV counts for the protein-only (Protein$^+$& miRNA$^-$), the miRNA-only (Protein$^-$& miRNA$^+$), and the dual-marker (Protein$^+$&miRNA$^+$) EVs among the total EVs detected. A total of 20 protein/miRNA combinations were examined, and 10 images were taken, with the average counts and standard deviations reported in the plots. d-g) The density distribution profiles of the fluorescence intensity ratios of all individual particles detected in the images for the four miRNA markers: d) -- let-7a; e) -- miR-21; f) -- miR-122; and g) -- miR-155. The ratio was calculated by dividing the miRNA signal with the sum intensity of the both the protein and miRNA signals.

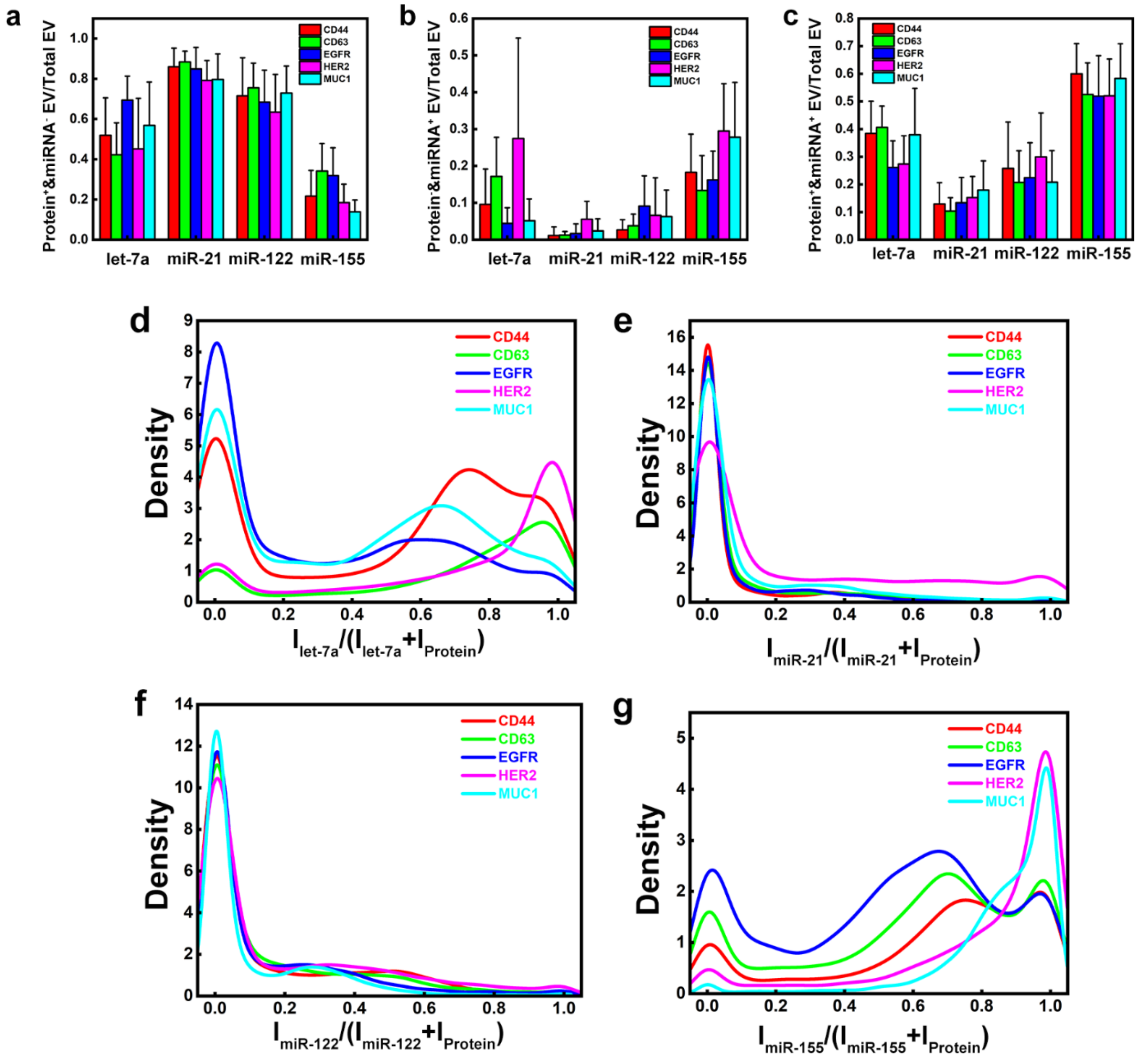


**Figure S16. Results of dual-marker NOBEL-SPA on the MCF-7 -derived EVs**. a-c) The bar plots of the fractions of the EV counts for the protein-only (Protein$^{+}$& miRNA$^{-}$), the miRNA-only (Protein$^{-}$& miRNA$^{+}$), and the dual-marker (Protein$^{+}$&miRNA$^{+}$) EVs among the total EVs detected. A total of 20 protein/miRNA combinations were examined, and 10 images were taken, with the average counts and standard deviations reported in the plots. d-g) The density distribution profiles of the fluorescence intensity ratios of all individual particles detected in the images for the four miRNA markers: d) -- let-7a; e) -- miR-21; f) -- miR-122; and g) -- miR-155. The ratio was calculated by dividing the miRNA signal with the sum intensity of the both the protein and miRNA signals.

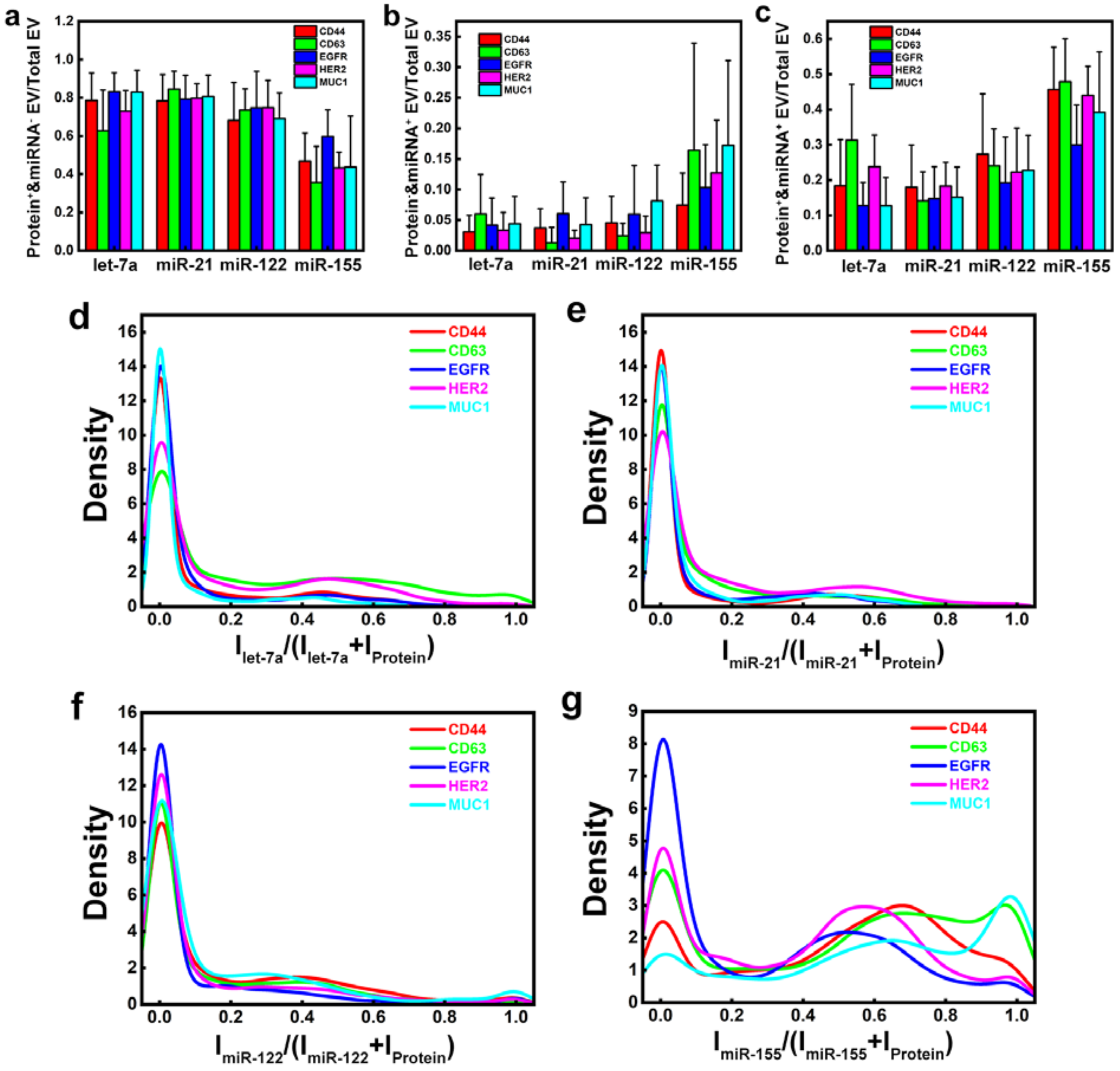


**Figure S17. Results of dual-marker NOBEL-SPA on the MDA-MB-231-derived EVs**. a-c) The bar plots of the fractions of the EV counts for the protein-only (Protein$^+$& miRNA$^-$), the miRNA-only (Protein$^-$& miRNA$^+$), and the dual-marker (Protein$^+$&miRNA$^+$) EVs among the total EVs detected. A total of 20 protein/miRNA combinations were examined, and 10 images were taken, with the average counts and standard deviations reported in the plots. d-g) The density distribution profiles of the fluorescence intensity ratios of all individual particles detected in the images for the four miRNA markers: d) -- let-7a; e) -- miR-21; f) -- miR-122; and g) -- miR-155. The ratio was calculated by dividing the miRNA signal with the sum intensity of the both the protein and miRNA signals.

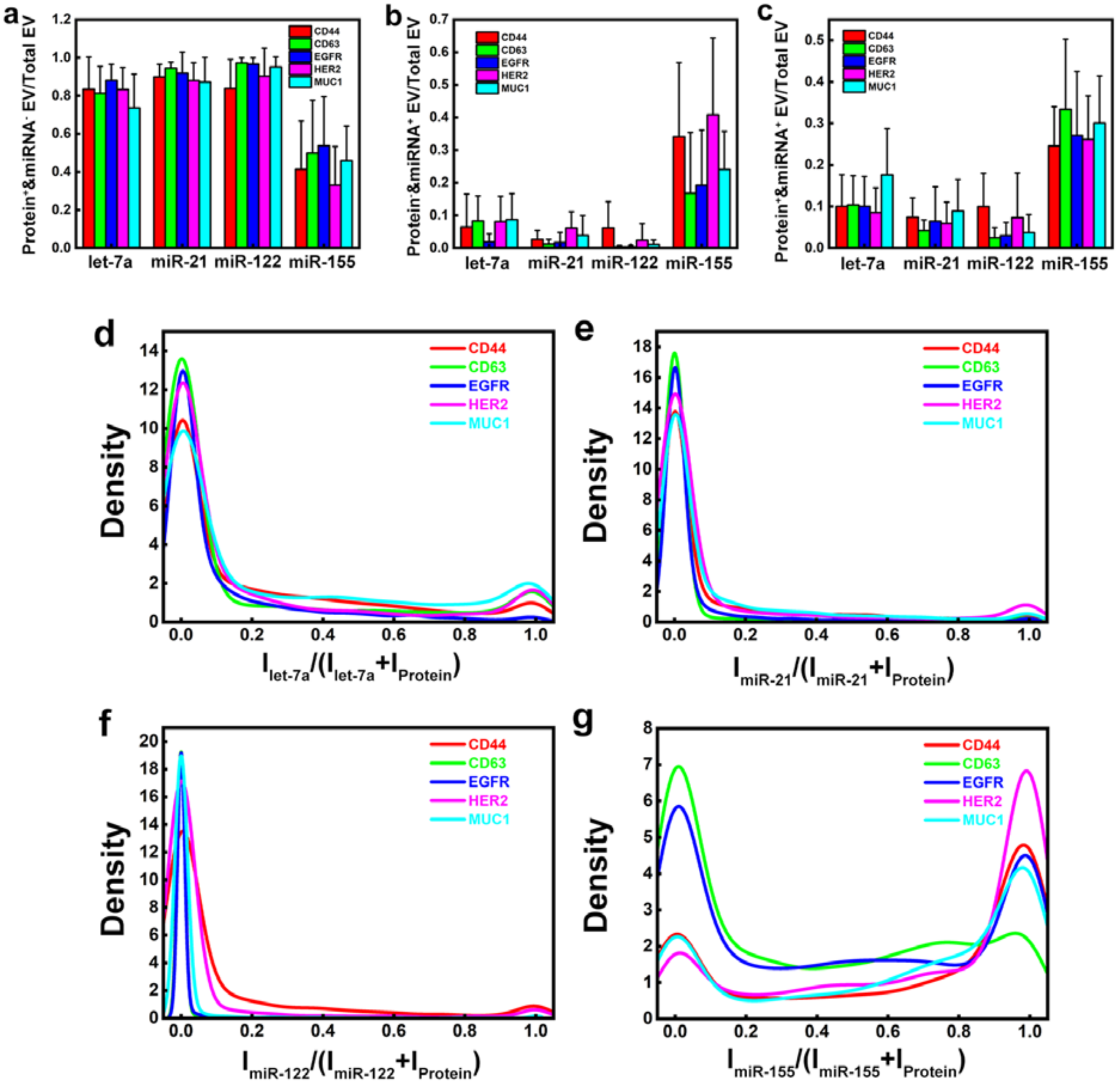


**Figure S18. Results of dual-marker NOBEL-SPA on the MCF-10A-derived EVs**. a-c) The bar plots of the fractions of the EV counts for the protein-only (Protein$^+$& miRNA$^-$), the miRNA-only (Protein$^-$& miRNA$^+$), and the dual-marker (Protein$^+$&miRNA$^+$) EVs among the total EVs detected. A total of 20 protein/miRNA combinations were examined, and 10 images were taken, with the average counts and standard deviations reported in the plots. d-g) The density distribution profiles of the fluorescence intensity ratios of all individual particles detected in the images for the four miRNA markers: d) -- let-7a; e) -- miR-21; f) -- miR-122; and g) -- miR-155. The ratio was calculated by dividing the miRNA signal with the sum intensity of the both the protein and miRNA signals.

**a**

| | Predicted Group | | | | | |
|---|---|---|---|---|---|---|
| | MDA-MB-231 | MCF-10A | A549 | HeLa | MCF-7 | Total |
| MDA-MB-231 | 20 | 0 | 0 | 0 | 0 | 20 |
| | 100.00% | 0.00% | 0.00% | 0.00% | 0.00% | 100.00% |
| MCF-10A | 0 | 20 | 0 | 0 | 0 | 20 |
| | 0.00% | 100.00% | 0.00% | 0.00% | 0.00% | 100.00% |
| A549 | 0 | 0 | 20 | 0 | 0 | 20 |
| | 0.00% | 0.00% | 100.00% | 0.00% | 0.00% | 100.00% |
| HeLa | 0 | 0 | 0 | 20 | 0 | 20 |
| | 0.00% | 0.00% | 0.00% | 100.00% | 0.00% | 100.00% |
| MCF-7 | 1 | 0 | 0 | 0 | 19 | 20 |
| | 5.00% | 0.00% | 0.00% | 0.00% | 95.00% | 100.00% |
| Total | 21 | 20 | 20 | 20 | 19 | 100 |
| | 21.00% | 20.00% | 20.00% | 20.00% | 19.00% | 100.00% |

**b**

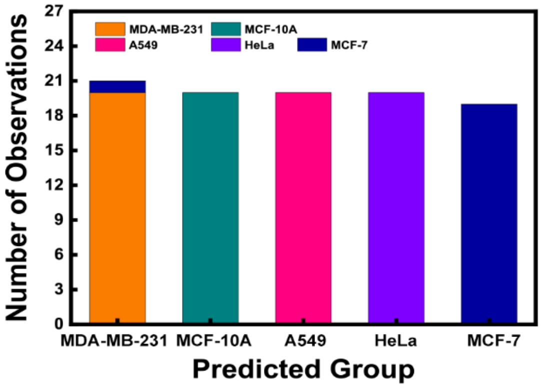


**Fig. S19. Canonical classification summary table (a) and plot (b) using the dual-marker counts for the 20 protein/miRNA combinations.**

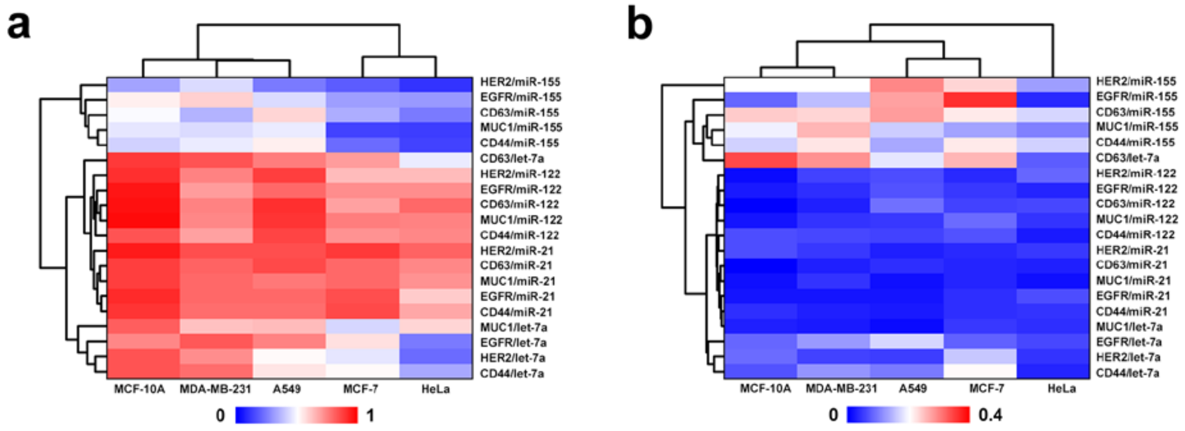


**Fig. S20. Heatmap of two EV sub-population: (a) protein$^+$/miRNA$^-$ EVs; and (b) protein$^-$/miRNA$^+$ EV.**

a

| | Predicted Group | | | | | |
|---|---|---|---|---|---|---|
| | MDA-MB-231 | MCF-10A | A549 | HeLa | MCF-7 | Total |
| MDA-MB-231 | 20 | 0 | 0 | 0 | 0 | 20 |
| | 100.00% | 0.00% | 0.00% | 0.00% | 0.00% | 100.00% |
| MCF-10A | 0 | 20 | 0 | 0 | 0 | 20 |
| | 0.00% | 100.00% | 0.00% | 0.00% | 0.00% | 100.00% |
| A549 | 0 | 0 | 20 | 0 | 0 | 20 |
| | 0.00% | 0.00% | 100.00% | 0.00% | 0.00% | 100.00% |
| HeLa | 0 | 0 | 0 | 19 | 1 | 20 |
| | 0.00% | 0.00% | 0.00% | 95.00% | 5.00% | 100.00% |
| MCF-7 | 0 | 0 | 0 | 0 | 20 | 20 |
| | 0.00% | 0.00% | 0.00% | 0.00% | 100.00% | 100.00% |
| Total | 20 | 20 | 20 | 19 | 21 | 100 |
| | 20.00% | 20.00% | 20.00% | 19.00% | 21.00% | 100.00% |

b

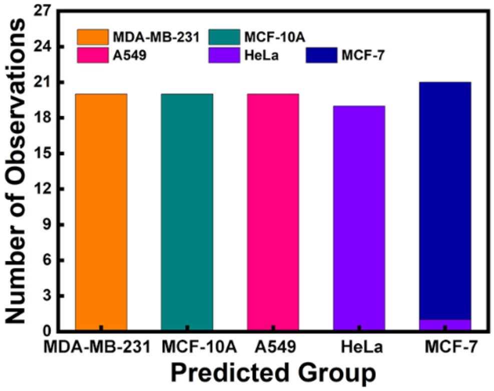


**Fig. S21. Canonical classification summary table (a) and plot (b) using the protein only counts for the 20 protein/miRNA combinations.**

**a**

| | Predicted Group | | | | | |
|---|---|---|---|---|---|---|
| | MDA-MB-231 | MCF-10A | A549 | HeLa | MCF-7 | Total |
| MDA-MB-231 | 19 | 0 | 0 | 0 | 1 | 20 |
| | 95.00% | 0.00% | 0.00% | 0.00% | 5.00% | 100.00% |
| MCF-10A | 0 | 18 | 1 | 0 | 1 | 20 |
| | 0.00% | 90.00% | 5.00% | 0.00% | 5.00% | 100.00% |
| A549 | 0 | 3 | 14 | 0 | 3 | 20 |
| | 0.00% | 15.00% | 70.00% | 0.00% | 15.00% | 100.00% |
| HeLa | 1 | 0 | 0 | 18 | 1 | 20 |
| | 5.00% | 0.00% | 0.00% | 90.00% | 5.00% | 100.00% |
| MCF-7 | 2 | 1 | 1 | 0 | 16 | 20 |
| | 10.00% | 5.00% | 5.00% | 0.00% | 80.00% | 100.00% |
| Total | 22 | 22 | 16 | 18 | 22 | 100 |
| | 22.00% | 22.00% | 16.00% | 18.00% | 22.00% | 100.00% |

**b**

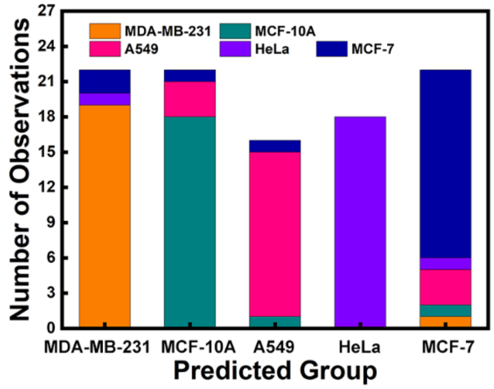


**C**

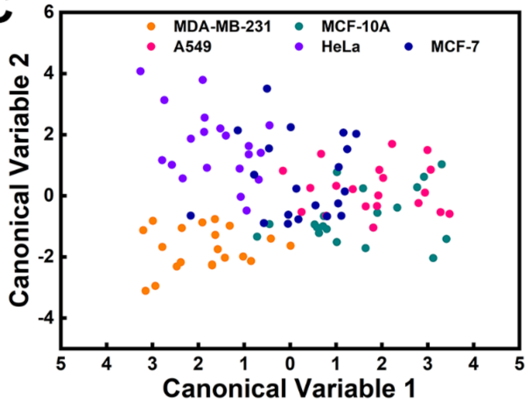


**Fig. S22. Canonical classification summary table (a), plot (b), and classification plot (c) using the RNA only counts for the 20 protein/miRNA combinations.**

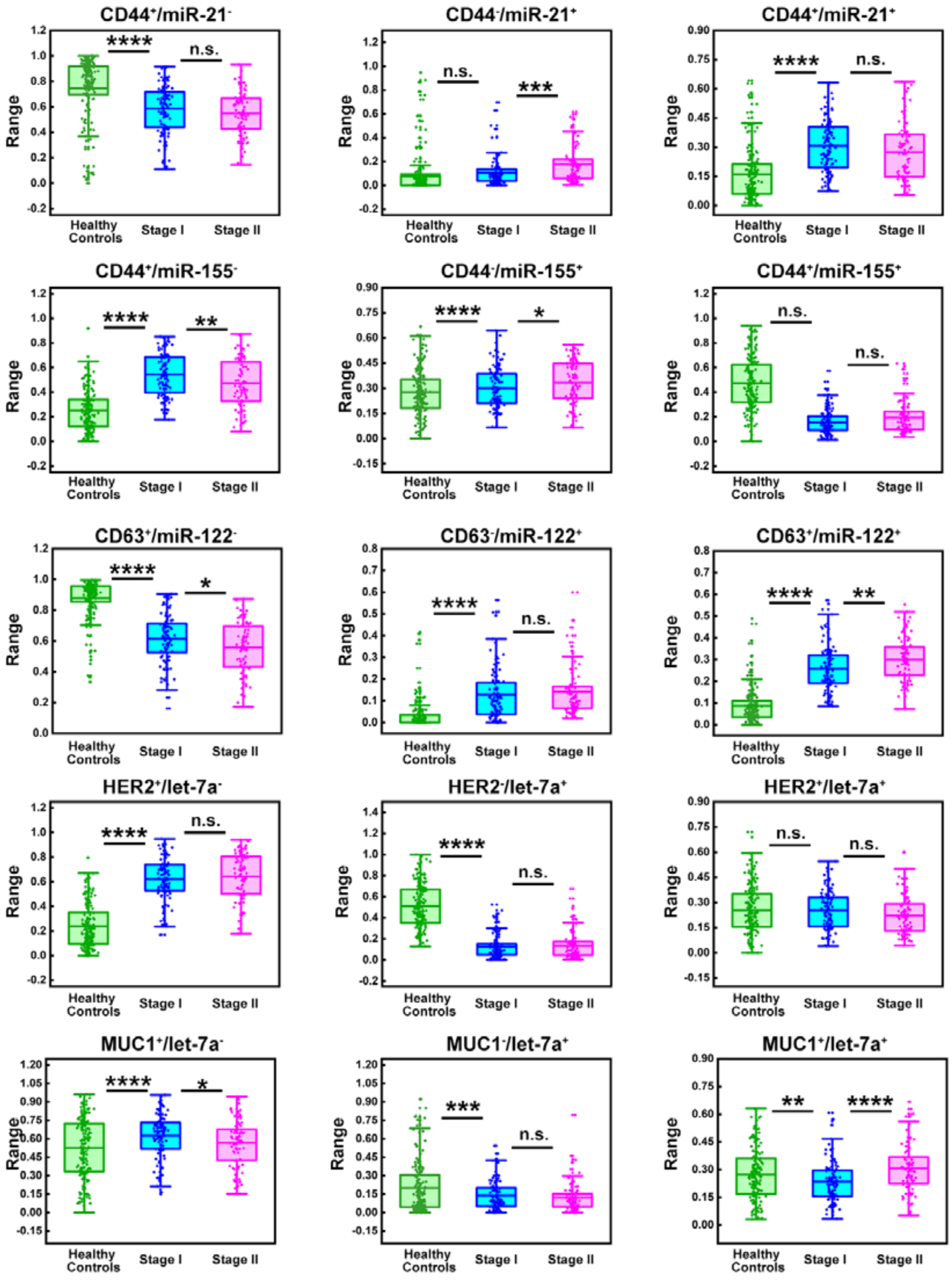


**Figure S23. Box plot of the proportions of the different exosome sub-populations (protein$^+$/miRNA$^-$, protein$^-$/miRNA$^+$, and protein$^+$/miRNA$^+$) among the total exosomes detected in sera samples collected from the healthy controls, stage I or II BC patients.**

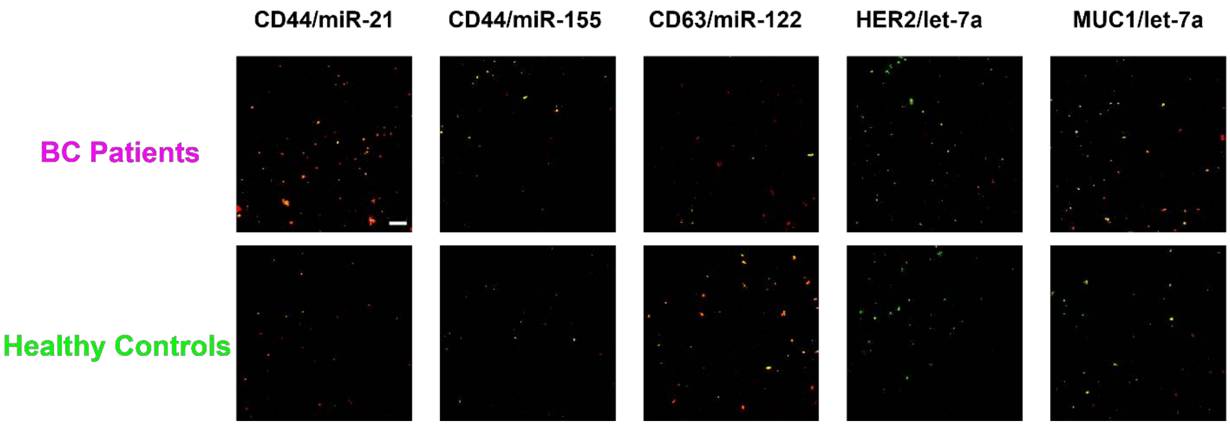


**Figure S24. Representative CFM images for dual-marker detection of selected 5 protein/miRNA combinations (CD44/miR-21. CD44/miR-155, CD63/miR-122, HER2/let-7a, MUC1/let-7a) in sera samples collected healthy controls and BC patients.** The protein$^{+}$/miRNA$^{-}$ EVs were marked in red; the protein$^{-}$/miRNA$^{+}$ EVs marked in green; and the protein$^{+}$/miRNA$^{+}$ EVs marked in orange. The scale bar was 10 µm.
